# Phylogenomics, Biogeography, and a New Family-level Classification of Silversides, Rainbowfishes, and Allies (Teleostei: Atheriniformes)

**DOI:** 10.64898/2026.05.05.722987

**Authors:** Lily C. Hughes, Victor de Brito, Kyle R. Piller, Seishi Kimura, Peter J. Unmack, Dahiana Arcila, Ricardo Betancur-R., Devin D. Bloom, Guillermo Ortí

**Affiliations:** Ichthyology Unit, North Carolina Museum of Natural Sciences, Raleigh, NC, USA 27601; Department of Marine, Earth, and Atmospheric Sciences, North Carolina State University, Raleigh, NC, USA 27607; Department of Biological Sciences, Western Michigan University, Kalamazoo, MI, USA 49008; Department of Biological Sciences, Southeastern Louisiana University, Hammond, LA, USA 70402; Fisheries Research Laboratory, Mie University, 4190-172 Wagu, Shima-cho, Shima, Mie, 517-0703, Japan; Centre for Applied Water Science, Institute of Applied Ecology, University of Canberra ACT 2617, Australia; Scripps Institution of Oceanography, University of California San Diego, 9500 Gilman Drive, La Jolla, CA 92093, USA; Department of Biological Sciences, George Washington University, Washington, DC, USA 20052; Department of Vertebrate Zoology, National Museum of Natural History, Smithsonian Institution, Washington, DC, USA

## Abstract

The order Atheriniformes (silversides, rainbowfishes, and blue-eyes) is a globally distributed group of fishes with frequent evolutionary transitions between marine and freshwater ecosystems. However, understanding the tempo and mode of these transitions has been hampered by poor phylogenetic resolution and limited taxonomic sampling, particularly within the suborder Atherinoidei. We generated a phylogenomic dataset of 1,100 exon loci for 150 species to resolve interfamilial relationships and reconstruct the group’s biogeographic history. We were also able to incorporate a large number of existing GenBank sequences, producing a phylogeny with 265 species sampled for at least some genetic data (67% of known species diversity). While the New World suborder Atherinopsidae is well-resolved, we found that the family Atherinidae is polyphyletic across all analyses. We propose a revised classification that restricts Atherinidae to the genus *Atherina* and recognizes Atherinomoridae and Craterocephalidae as separate families. Our biogeographic inferences using explicit geographic areas suggests more frequent marine-to-freshwater transitions than previously inferred with simplified binary (marine vs. freshwater) coding, and uncover habitat transitions where marine ancestors may have gone extinct. These results highlight how explicit geographic modeling can uncover marine ancestry erased by extinction, providing a robust phylogenetic framework for future evolutionary studies of Atheriniformes.

## 1. Introduction

The fish order Atheriniformes is globally distributed in temperate and tropical coastal marine waters and freshwater lakes and streams, and currently contains over 390 valid species (Fricke et al., 2026). Many members possess a distinctive silver lateral stripe, giving rise to their common name – silversides. Many species are relatively small-bodied, and form large schools that are ecologically important forage fish in marine environments. Atheriniformes have established independent freshwater radiations on multiple continents and islands (Bloom et al., 2013; Hughes et al., 2020; Stelbrink et al., 2014; Unmack et al., 2013). This makes them an excellent study system for studying phenotypic and ecological evolution associated with transitions between marine and freshwater ecosystems (de Brito et al., 2022), if a robust phylogeny can be established.

Atheriniformes have a long history of support from morphological evidence as members of Atherinomorpha – which also includes Beloniformes, and Cyprinodontiformes (Parenti, 1993). The eleven families are divided into two suborders, with the Atherinopsidae “New World” silversides forming the suborder Atherinopsoidei, and all other families falling into Atherinioidei, which is supported by both molecular and morphological evidence (Bloom et al., 2012; Campanella et al., 2015; Dyer and Chernoff, 1996). Early morphological studies relegated most families to a polytomy of atherinoids (Stiassny, 1990). Family-level classifications have changed with the use of molecular studies, for example supporting the placement of *Notocheirus* within Atherinopsidae, instead of with the genus *Iso* (Bloom et al., 2012).

However, the relationships of several families are still poorly resolved by both morphological and molecular datasets. For example, the largest molecular dataset analyzed to date used eight loci (one mitochondrial and seven nuclear loci) and found Pseudomugilidae and Telmatherinidae were nested within Melanotaeniidae, but there was no statistical support for these relationships, effectively leaving these relationships unresolved (Campanella et al., 2015).

Long-branch attraction (LBA) is a long-known bias in phylogenetics (Felsenstein, 1978; Philippe et al., 2005; Susko and Roger, 2021), has also hindered the systematics of Atheriniformes. In particular, the melanotaeniid genus *Cairnsichthys* sometimes breaks away from the rest of the family (Aarn and Ivantsoff, 1997; Bloom et al., 2012; Campanella et al., 2015). Families Atherionidae and Phallostethidae have long branches that sometimes attract towards outgroups in phylogenetic analyses (Sparks and Smith, 2004), or towards each other (Campanella et al., 2015). Recently, more complex substitution models accounting for heterotachy have been developed (Crotty et al., 2019), and have been shown to ameliorate persistent LBA artifacts in flatfishes (Duarte-Ribeiro et al., 2024). Outgroup selection also plays a role in LBA artifacts, particularly where outgroups are distantly related to the in-group (DeSalle et al., 2023; Li et al., 2012; Simmons et al., 2022). More fully accounting for LBA in the position of long-branched lineages in Atheriniformes could help improve the resolution of interfamilial relationships.

A lack of clarity of atheriniform relationships hampers our biogeographic interpretation of this widespread group. While many fish lineages are stenohaline and can only tolerate narrow salinity ranges, Atheriniformes are notable for their repeated transitions between marine and freshwater biomes, with many species frequenting estuaries (Bamber and Henderson, 1988).

Marine-freshwater transitions are frequently modeled as binary states to examine habitat evolution and diversification both within Atheriniformes (Bloom et al., 2013; Campanella et al., 2015) and across the fish Tree of Life (Betancur-R et al., 2015; Rabosky, 2020). However, this simplified binary coding masks the geographical complexity of their distributions and could underestimate the true number of habitat transitions. Several large atheriniform freshwater radiations on the Sahul shelf (present-day Australia and New Guinea) and Madagascar have no living marine relatives, and their distribution was previously hypothesized to be the result of Gondwanan vicariance (Sparks and Smith, 2004). But large scale studies of ray-finned fishes indicate that the origin of Atheriniformes is in the Late Cretaceous (Betancur-R. et al., 2013; Ghezelayagh et al., 2022; Hughes et al., 2018; Near et al., 2013), after the breakup of the southern continents, making marine dispersal the most plausible origin for these freshwater radiations. Explicit coding of biogeographical areas, rather than a two-state binary character, should provide a more realistic understanding of the number of habitat shifts in Atheriniformes.

Phylogenomic-scale datasets comprising hundreds of genes are now accessible across a wide range of taxonomic groups. We sequenced a set of 1,100 single-copy, orthologous exon loci using a target-capture probe set designed for the fish clade Ovalentaria (Hughes et al., 2021) for all atheriniform families except the monotypic Dentatherinidae, which has never been sequenced. We inferred a phylogeny for Atheriniformes from this genome-scale dataset using a variety of analyses, including the use of heterotachy models and long branch-exclusion for long-branching taxa in Atherinoidei. We incorporated GenBank data to increase taxonomic sampling, and estimated a time-calibrated phylogeny of 265 atheriniform species – 67% of known species diversity in this order, and more than doubling the taxonomic sampling of 103 species in Campanella *et al*. (2015). Finally, we conducted a biogeographical analysis using the DEC model, accounting for multiple marine and freshwater areas across the globe.

## 2. Methods

### 2.1 Sequencing & Exon Assembly

DNA was extracted from 150 Atheriniformes tissue samples (149 species) and seven outgroup tissues from Beloniformes on a Autogen 96-well plate platform at the Laboratory of Analytical Biology at the Smithsonian Institution (Washington, DC). Library preparation and sequence capture was conducted by Arbor Biosciences (Ann Arbor, MI), using the Ovalentaria exon probe set developed by Hughes *et al*. (2021). Illumina libraries were sequenced as 100 bp paired-end reads on a HiSeq 4000 at the University of Chicago. Raw sequencing reads were processed and assembled into loci using the pipeline of Hughes *et al*. (2021). Raw reads were trimmed using Trimmomatic v.0.39 (Bolger et al., 2014) for low quality bases and adapter contamination. Reads were mapped against percomorph reference sequences with BWA v.0.7.17 (Li and Durbin, 2009), and an initial sequence assembled for each locus with Velvet v.1.2.10 (Zerbino and Birney, 2008), then used as a reference sequence for iterative locus assembly with aTRAM 2.0 (Allen et al., 2018) with the Trinity v.2.2 assembler (Grabherr et al., 2011). Identical contigs were removed with CD-Hit-EST v.4.8.1 (Fu et al., 2012), then open reading frames (ORFs) were identified with Exonerate v.2.4.0 (Slater and Birney, 2005). If multiple contigs with ORFs and over 1% sequence divergence were assembled, they were excluded from downstream analyses. To verify species identities, we searched assembled COI sequences in the Barcode of Life Database (BOLD; Ratnasingham and Hebert, 2007), though not all atheriniforms are represented in the BOLD database. We incorporated sequences from additional published genomes and transcriptomes into our final matrix (Hughes et al., 2018; Vega-Retter et al., 2018; Wilder et al., 2020). Accession numbers and voucher specimens are listed in Table S1, and raw data can be accessed at NCBI BioProject XXXXXX (pending).

### 2.2 Matrix Assembly & Phylogenomic Analysis

We assembled multiple matrices with differing amounts of missing data: a “full” data set containing 1,089 loci, a “G75” data set containing 735 loci with at least 75% present data, and a “G90” data set with 420 loci with at least 90% present data. Each of these matrices with different amounts of missing data were also translated to amino acids. These matrices were analyzed under maximum likelihood (ML) using IQ-TREE 2.2.0 as both nucleotides and amino acids (Minh et al., 2020). Nucleotide matrices were divided into three codon positions and assigned separate GTR+G4 models. The amino acid data set was not partitioned, and the best model was selected using ModelFinder (Kalyaanamoorthy et al., 2017). Support was assessed with 1,000 ultrafast bootstrap replicates (Hoang et al., 2018). To examine the position of three long-branched families (Atherionidae, Isonidae, and Phallostethidae), we also created matrices that excluded each family. We tested for the effect of potential outgroup attraction of these three families by reducing the number of outgroups to two members of Atherinopsidae, and analyzed the full, G75, and G90 gene sets with this reduced taxon sampling. We also analyzed a taxon-reduced G75 dataset under the GHOST model, which models heterotachy (Crotty et al., 2019).

For multispecies coalescent (MSC) analysis, we estimated individual gene trees with IQ-TREE, also with 1,000 ultrafast bootstrap replicates. Nodes with less than 20% support were collapsed before input into ASTRAL-III (Zhang et al., 2018). We analyzed the full, G75, and G90 nucleotide gene sets in ASTRAL, as well as the same datasets with reduced outgroup sampling. A full list of analyses and dataset sizes is included in Table 1.

**Table 1.** Summary of phylogenomic analyses performed in this study. Concatenated Maximum Likelihood analyses were conducted in nucleotide and amino acid matrices at the three levels of gene completeness (all loci, 75% complete, and 90% complete). Taxa, number of loci, total alignment length (in base pairs for nucleotide or amino acids matrices), internal analysis name, and analysis number are provided for each dataset.

| Description | Taxa | Loci | Sites | Analysis Name | Analysis Number |
| --- | --- | --- | --- | --- | --- |
| **Concatenation** |  |  |  |  |  |
| All Nucleotides | 158 | 1,089 | 443,706 bp | full.nt | 1 |
| G75 Nucleotides - 75% complete matrix | 158 | 735 | 333,915 bp | g75.nt | 2 |
| G90 Nucleotides - 90% complete matrix | 158 | 420 | 196,569 bp | g90.nt | 3 |
| All Amino Acids | 158 | 1,089 | 147,902 aa | full.aa | 4 |
| G75 Amino Acids | 158 | 735 | 111,305 aa | g75.aa | 5 |
| G90 Amino Acids | 158 | 420 | 65,523 | g90.aa | 6 |
| **Long Branch Exclusion** |  |  |  |  |  |
| Atherionidae Excluded | 72 | 1,089 | 443,706 bp | no_atherion | 7 |
| Isonidae Excluded | 71 | 1,089 | 443,706 bp | no_iso | 8 |
| Phallostethidae Excluded | 68 | 1,089 | 443,706 bp | no_phallo | 9 |
| All Loci - Reduced Outgroups | 73 | 1,039 | 432,141 bp | all_redout | 10 |
| G75 - Reduced Outgroups | 73 | 726 | 329,601 bp | g75_redout | 11 |
| G90 - Reduced Outgroups | 73 | 455 | 214,503 bp | g90_redout | 12 |
| **Heterotachy** |  |  |  |  |  |
| GHOST model - G75 dataset with reduced taxon sampling | 50 | 735 | 329,601 bp | GHOST | 13 |
| **Multispecies Coalescent** |  |  |  |  |  |
| All Gene Trees | 158 | 1,089 |  | full.c20 | 14 |
| G75 Gene Trees | 158 | 735 |  | g75.c20 | 15 |
| G90 Gene Trees | 158 | 420 |  | g90.c20 | 16 |
| All Gene Trees - Reduced Outgroups | 73 | 1,039 |  | full.c20.redout | 17 |
| G75 Gene Trees - Reduced Outgroups | 73 | 726 |  | g75.c20.redout | 18 |
| G90 Gene Trees - Reduced Outgroups | 73 | 455 |  | g90.c20.redout | 19 |

### 2.3 Extending Taxonomic Sampling with GenBank Data

Our exon capture probe set was designed to include a number of loci that have long standing use in fish phylogenetics (Hughes et al., 2021). To maximize taxonomic sampling for biogeographic and comparative analysis, we incorporated GenBank sequences from previously published studies. These genes included mitochondrial loci: COI, CytB, ND1, ND2, and ND5; as well as nuclear sequences: FICD, GCS1, KBTBD4, KIAA1239, MYH6, PANX2, RAG1, SH2PX2, and SLC10A3. Tree topologies inferred with the G75 nucleotide matrix under both ML and MSC methods were each used as a backbone constraint where the topology of taxa with phylogenomic data was kept constant. Taxa with GenBank-only data were constrained to families. The GenBank sequences used are listed in Table S2, and come from a wide range of studies at different taxonomic levels (Allen et al., 2016, 2015; Gerald R. Allen et al., 2014; Gerald R Allen et al., 2014; Allen and Unmack, 2012; Bloom et al., 2013, 2012, 2009; Campanella et al., 2015; Hughes et al., 2020; Kadarusman et al., 2012; Lau and Jacobs, 2017; Page and Hughes, 2010; Sasaki and Kimura, 2020, 2014; Setiamarga et al., 2008; Sparks and Smith, 2004; Unmack et al., 2013; Unmack and Dowling, 2010; Valdez-Moreno et al., 2009; Xu et al., 2021).

### 2.4 Divergence Time Estimation

To address computational challenges, we used a reduced data subset containing the top twelve exon loci with the least missing data as well as all nuclear and mitochondrial loci mined from GenBank above. The reduced matrix had an alignment length of 19,797 bp. We obtained the best fitting model with ModelFinder (Kalyaanamoorthy et al., 2017) for each of three codon positions and set those models in BEAUti (Bouckaert et al., 2019). We used the Optimized Relaxed Clock model in BEAST 2.7.6 (Bouckaert et al., 2019; Douglas et al., 2021). We used the two topologies obtained from the previous analyses including GenBank data, one based on the G75 ML analysis and the other on the G75 MSC analysis, and conducted two separate searches constraining those topologies. Two independent runs for each topology were conducted for 400,000,000 generations. Atheriniform fossils are scarce, but we assigned four calibrations on nodes in our tree. We used RelTime (Tamura et al., 2018) to generate initial ultrametric starting trees that satisfied these calibration points.

1. Root Atheriniformes: †*Rhamphognathus paralepoides*, **MNHN Bol 10 (10874)**. This fossil belongs to an extinct atherinoid family from the Monte Bolca Lagerstätten (Bannikov, 2008), which dates to 49.1 Ma, and provides the minimum age for the group. The soft upper bound was determined from the literature. Large scale phylogenetic analyses of fishes have consistently placed the origin of Atheriniformes to the latest Cretaceous, from 70-75 Ma (Betancur-R. et al., 2013; Ghezelayagh et al., 2022; Hughes et al., 2018; Near et al., 2013). We used 75 Ma as the soft 95% maximum bound.
2. Crown Atherinoidei: †*Hemitrichas stapfi* **SSN12DX12**, Heimat Museum, Germany. This fossil atherinid from the Mainz Basin dates to the Late Oligocene (23.02 Ma) based on stratigraphy (Gaudant and Reichenbacher, 2005). However, Atherinidae *sensu lato* was not monophyletic in any of our analyses; we therefore consider this fossil to be *incertae sedis* within Atherinoidei. Soft upper 95% bound was set to 49.1, based on the next oldest fossil.
3. Crown *Atherina*: *Atherina* †*atropatiensis* , **BSPG 2010 XXI−4** (Carnevale *et al*. 2011) (Carnevale et al., 2011). This fossil is closely related to extant *A. boyeri* based on meristic and morphological features. It was recovered from the lignite beds of the Tabriz Basin, NW Iran, which are dated to 11.6-10 Ma (Reichenbacher et al., 2011). We used 10 Ma as the minimum age for the MRCA of *Atherina boyeri* and *A. hepsetus*. The soft maximum age was set to 23.02 Ma based on the next oldest fossil
4. Crown *Odontesthes*: *Odontesthses sp.* **MLP 04-V-2-385** (Bogan et al., 2009). This fossil is of Middle Pleistocene age (0.23 Ma +/- 0.03) based on the 40Ar/39Ar age of impact glass bounding the upper limit of the fossil fish bearing layer at Centinela del Mar locality in Argentina (Schultz et al., 2004). Its relationship to *Odontesthes* species in the area is unclear; we therefore used it as a minimum age constraint for the entire La Plata species-group that now inhabits the Pampas region (MRCA *Odontesthes argentinensis* and *O. bonariensis*).

### 2.5 Biogeographic Analysis under the DEC Model

Biogeographic analysis under the best-fitting Dispersal-Extinction-Cladogenesis model (Ree and Smith, 2008) was executed in BioGeoBears (Matzke, 2018). We did not include the “founder event speciation” parameter *j*, as this is unlikely in our system (Ree and Sanmartín, 2018). We analyzed the two suborders separately as Atherinopsidae is restricted to the Americas and the remaining atherinoid families are distributed globally, though most species are restricted regionally. We considered nine areas for Atherinopsidae: Northwest Atlantic, North Atlantic Freshwater, Tropical Western Atlantic, Tropical Atlantic Freshwater, Southwest Atlantic, Southwest Atlantic Freshwater, Southeast Pacific, Southeast Pacific Freshwater, Tropical Eastern Pacific, Tropical Pacific Freshwater, Northeast Pacific. Though they may ascend estuaries, no atherinopsids currently inhabit temperate Pacific-draining freshwater habitats, so this area was not included. Species that primarily inhabit marine waters but inhabit estuaries were coded as marine; species that primarily inhabit freshwater but sometimes enter estuaries were coded for the appropriate freshwater habitat. Species recorded from both marine and freshwater records were assigned both habitats. Records from GBIF (limited to preserved specimens collected after 1950) and information from FishBase were used to categorize the distribution species (Froese and Pauly, 2025; “GBIF Occurrence Download,” 2024). Dispersal was only allowed between adjacent areas, and the maximum number of areas allowed was limited to three (this is the largest number of areas any living species inhabits). Our time-stratified analysis allowed dispersal between the tropical Atlantic and Pacific oceans until the final closure of the Isthmus of Panama at 2.8 Ma (O’Dea et al., 2016).

Biogeographical area coding for Atherinoidei consisted of six marine and four freshwater areas. The six marine areas were delineated from the Marine Ecoregions of the World: Western Indian Ocean, Indo-Pacific Ocean, Central Pacific Ocean, Temperate Australia, Western Atlantic, and Eastern Atlantic. The four freshwater areas considered were: Madagascar, Australia-New Guinea (Sahul), Sundaland, and Sulawesi. Located in Wallacea, Sulawesi has a complex history and is a composite island made by the collision of the Asian and Australian plates. The maximum number of areas allowed was three, the largest of any living atherinoid species.

The time stratified analysis had five different time slices based on geological events. Dispersal between freshwater Sundaland and Sulawesi was allowed before the opening of the deep Makassar Strait at approximately 40 Ma (Hall, 2012); after this time dispersal would have to occur through the Indo-Pacific Ocean. Approximately 20 million years ago, part of the Sahul shelf known as the Sula Spur collided with North Sulawesi (Hall, 2012) and created an opportunity for dispersal between Australia-New Guinea and Sulawesi freshwater habitats. This connection closed at 5 Ma, when eastern Sulawesi again became fully isolated from the Sahul shelf. Before the final closure of the Tethys at approximately 12 Ma, dispersal was allowed between the Eastern Atlantic and the Indo-Pacific Oceans through this seaway. Finally, after 2.8 Ma, dispersal from the Pacific to the Western Atlantic was blocked by the rise of the Isthmus of Panama. We also considered another potential biogeographic barrier: the Eastern Pacific Barrier (EPB), a 5,000-8,000 km stretch of deep water that separates the Eastern Pacific from the Central Pacific (Briggs, 1961). We used two different dispersal matrices to account for this barrier. In one scenario, the EPB was considered a low barrier to dispersal, allowing dispersal from the Central Pacific to the Western Atlantic until the closure of Panama. The other kept the dispersal probability across the EPB low (0.001), making the EPB a high barrier to dispersal and forcing any colonization of the Western Atlantic through the Eastern Atlantic instead. Beyond these time slices and dispersal probabilities, dispersal was only allowed between adjacent areas.

## 3. Results

### 3.1 Phylogenomics of Atheriniformes

Our full phylogenomic dataset had an alignment length of 443,706 bp. Atherinopsidae is consistently obtained as the sister group to all other families, and has long been recognized as a separate suborder. Relationships within Atherinopsidae are highly congruent and well-supported across analyses, with some low support associated with the position of *Melanorhinus microps* (Fig. 1). *Notocheirus hubbsi* is nested within this family, consistent with other molecular analyses, though morphological classifications placed it as the sister group to the genus *Iso*.

**Fig. 1.**
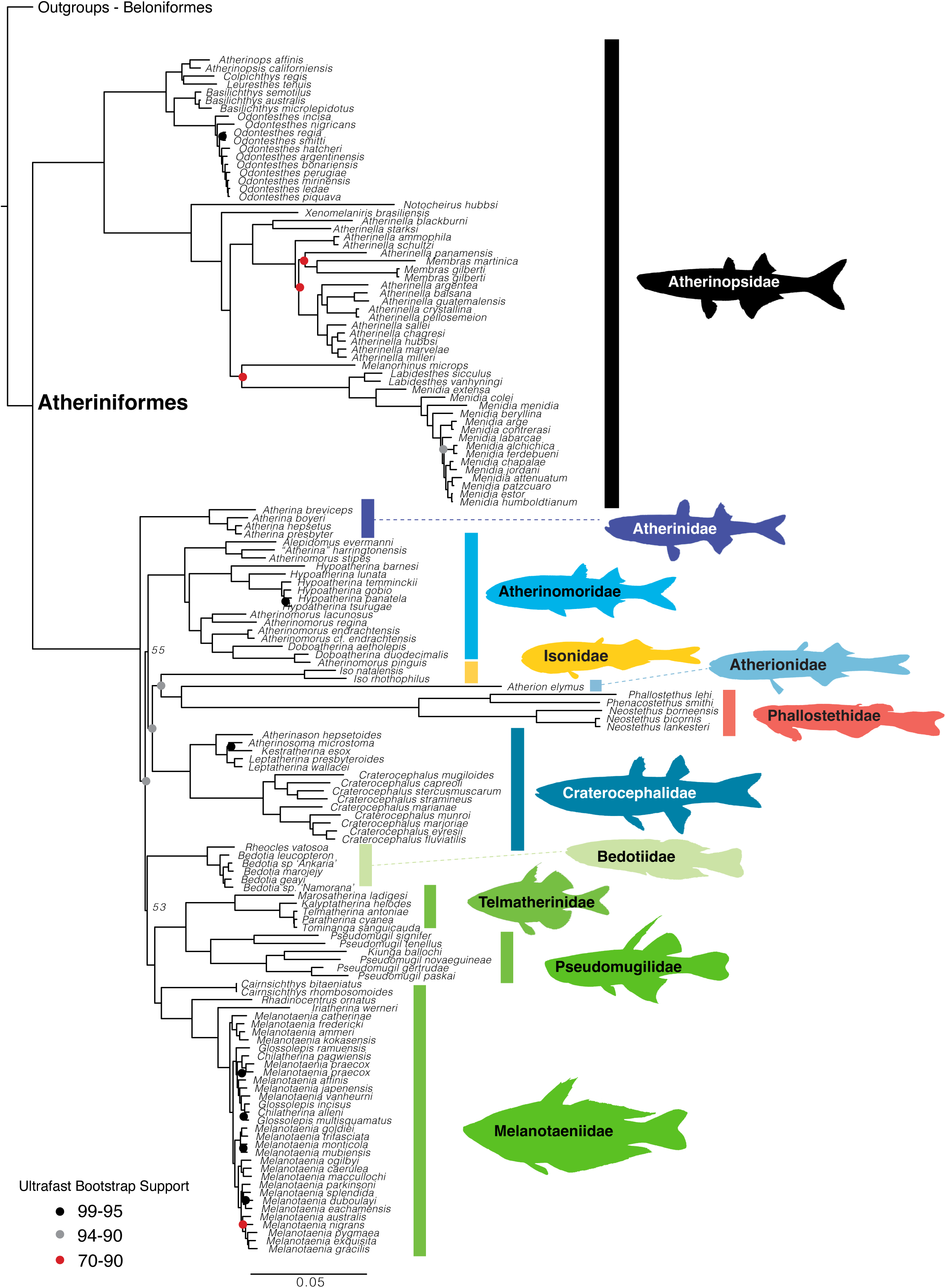
Phylogenomic tree of Atheriniformes based on the concatenated nucleotide G75 matrix analyzed in IQ-TREE (Table 1; Analysis 2). Nodes without annotations have 100% ultrafast bootstrap support. The tree was rooted with ten outgroup taxa from Beloniformes, which are not shown in the figure.

Relationships among the atherinoid families are more poorly supported and variable between ML and MSC analyses (Fig 2). This area of the tree is characterized by very short internodes at the base of the clade. Melanotaeniidae, Pseudomugilidae, and Telmatherinidae are supported as a clade across all analyses. In ML analyses, the long-branched families Atherionidae, Isonidae, and Phallostethidae tend to group together with relatively high support. Excluding these long branches does not change the position of the other branches in the ML trees. MSC analyses instead tended to favor Atherionidae and Phallostethidae as the first two branches of Atherinoidei (Fig. 2f-i).

**Fig. 2.**
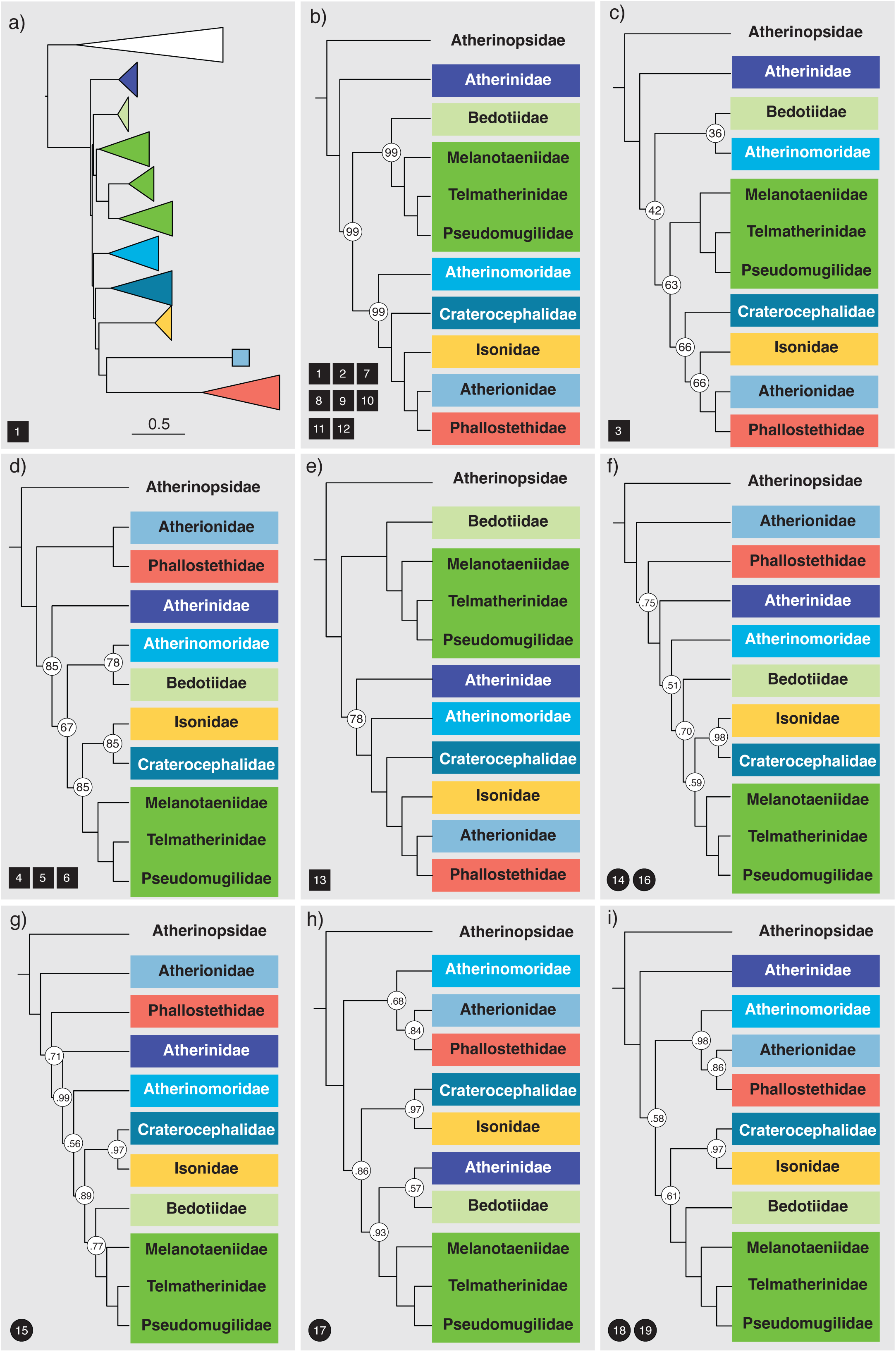
Comparison of conflicting family-level relationships in Atheriniformes across analyses. Numbers in black boxes or circles indicate the analysis number in Table 1. Black boxes indicate Maximum Likelihood (ML) results from IQ-TREE, and black circles indicate multispecies coalescent results from ASTRAL. Node support values reflect the first analysis listed when a given topology was obtained multiple times; nodes without support values received maximum support. a) Family level topology for Analysis 1, showing short internodal branch lengths among atherinoid families. b) Most common topology obtained in ML analyses of nucleotides for (Analyses 1, 2, 7-12). c) Topology obtained in Analysis 3 with the G90 matrix. d) Topology obtained from the analyses of amino acids (Analyses 4-6). e) Topology estimated under the GHOST model accounting for heterotachy (Analysis 13). f-i) Topologies obtained from ASTRAL analyses.

Across all 19 analyses, the family Atherinidae was never found monophyletic (Fig. 2; Figs. S1-S19). Instead, three separate and highly supported groups emerged: a group containing only the genus *Atherina* (=Atherinidae in Fig. 1); a group containing the Australian genus *Craterocephalus* sister to the Temperate Australian marine genera *Atherinason*, *Atherinasoma*, *Kestratherina*, and *Lepthatherina* (= Craterocephalidae in Fig. 1); the last group contains the genera *Alepidomus*, *Atherinomorus*, *Doboatherina*, and *Hypoatherina*, as well as the Western Atlantic species *“Atherina” harringtonensis* (=Atherinomoridae in Fig. 1).

### 3.2 Divergence Time Estimation

The earliest atherinform split was estimated to be in the latest Cretaceous, separating Atherinopsidae from the other families (Fig. 3a). Topological uncertainty did not significantly change divergence times, and both the ML and MSC-dated trees show median ages of initial diversification of both atherinopsids and atherinoids in the early Eocene (Fig. 3a-b).. Lineage accumulation is not steady through time, diverging significantly from Yule-model expectations. In Atherinopsidae, early lineages accumulated rather slowly, with recent freshwater radiations in *Menidia* and *Odontesthes* (Fig. 3c). Atherinoid lineages diverged rapidly in the Eocene, and all family-level lineages were established by the onset of the Oligocene (Fig. 4).

**Fig. 3.**
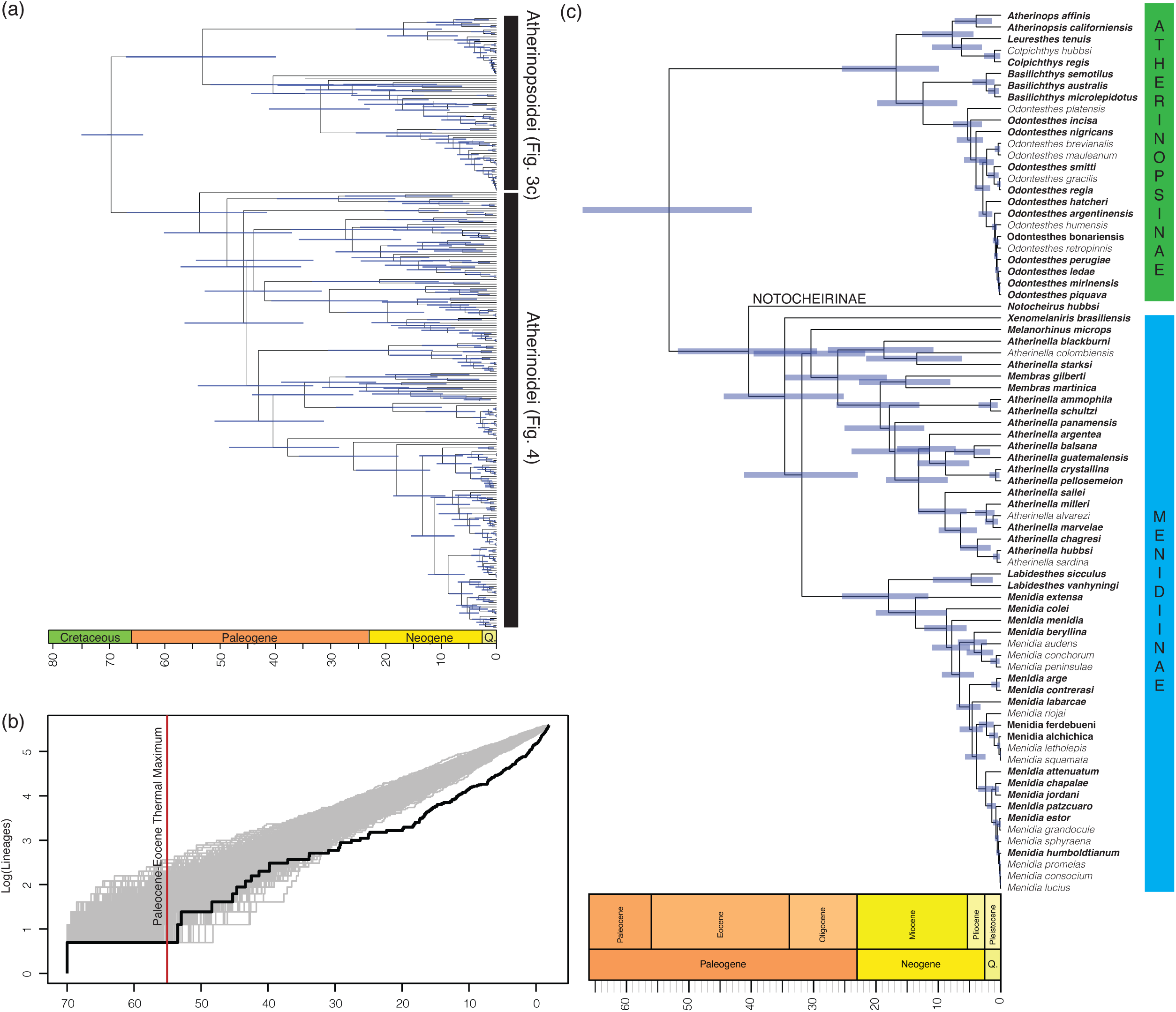
a) Time-calibrated phylogeny of Atheriniformes based on the ASTRAL topology, with taxa from GenBank sequences added to the phylogenomic backbone. Detailed phylogenies with taxon labels are shown in panel c and Fig. 4. b) Lineage-through-time plot of Atheriniformes. The black line indicates observed lineage accumulation, and gray lines represent simulations under a Yule model. c) Timescale of Atherinopsidae evolution. Bolded names indicate taxa supported by phylogenomic data, remaining taxa have only GenBank sequences. Blue node bars indicate 95% HPD intervals.

**Fig. 4.**
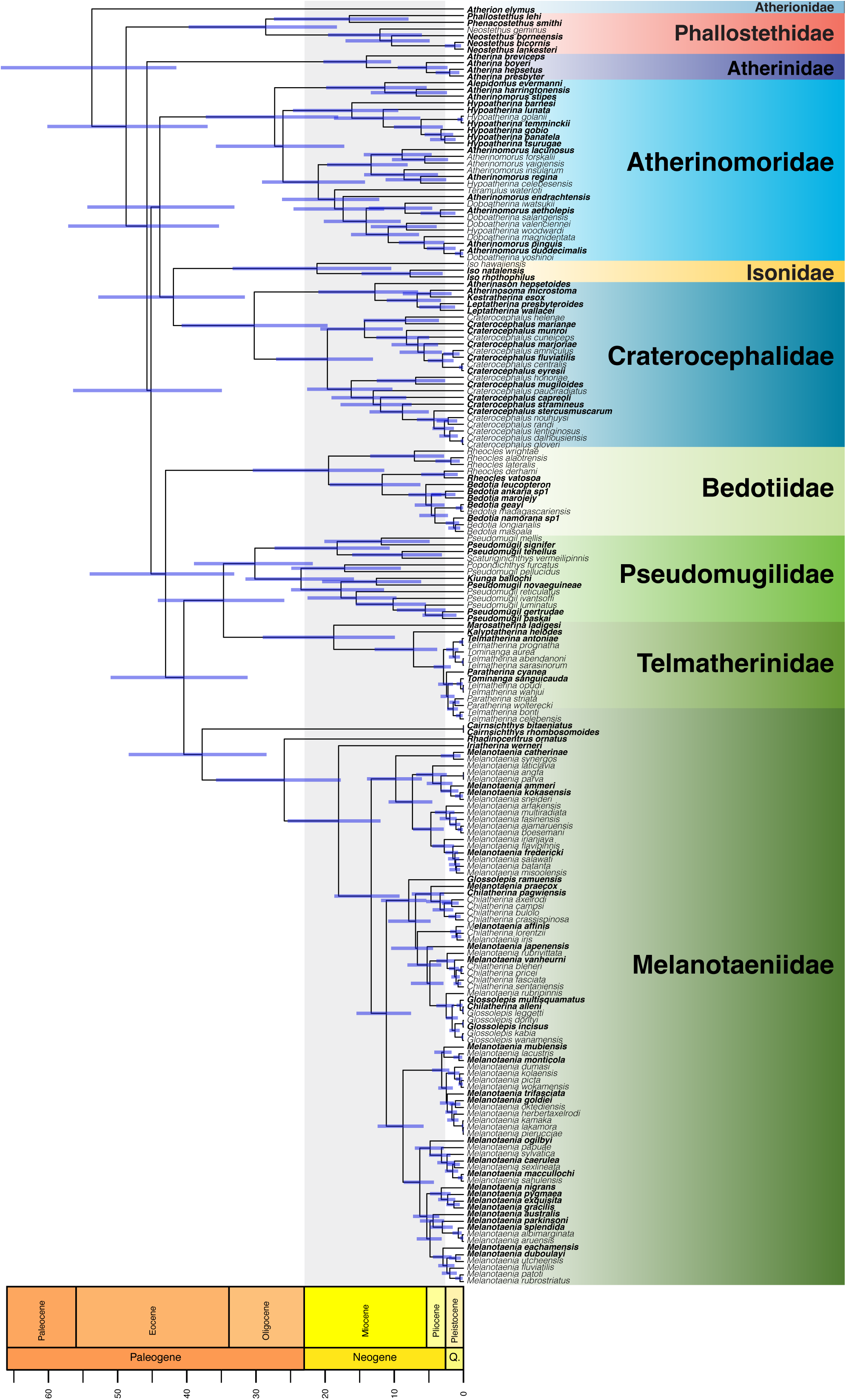
Timescale of atherinoid evolution based on the ASTRAL topology, and the addition of taxa from GenBank sequences. Bolded names indicate taxa supported by phylogenomic data, remaining taxa are represented by GenBank sequences only. Blue node bars indicate 95% HPD intervals.

### 3.3 Biogeographic Inference

Biogeographic analysis under the DEC model for Atherinopsidae’s history in the Americas suggests up to ten instances of freshwater colonizations by marine ancestors when considering their spread across eleven different regions (Fig. 5; Figs. S20-S23). Most ancestral nodes are reconstructed as marine, particularly in tropical waters in the Western Central Atlantic and Tropical Eastern Pacific. While the closure of the Isthmus of Panama 2.8 million years ago separated many sister species in other clades, the only sister species split between the Western

**Fig. 5.**
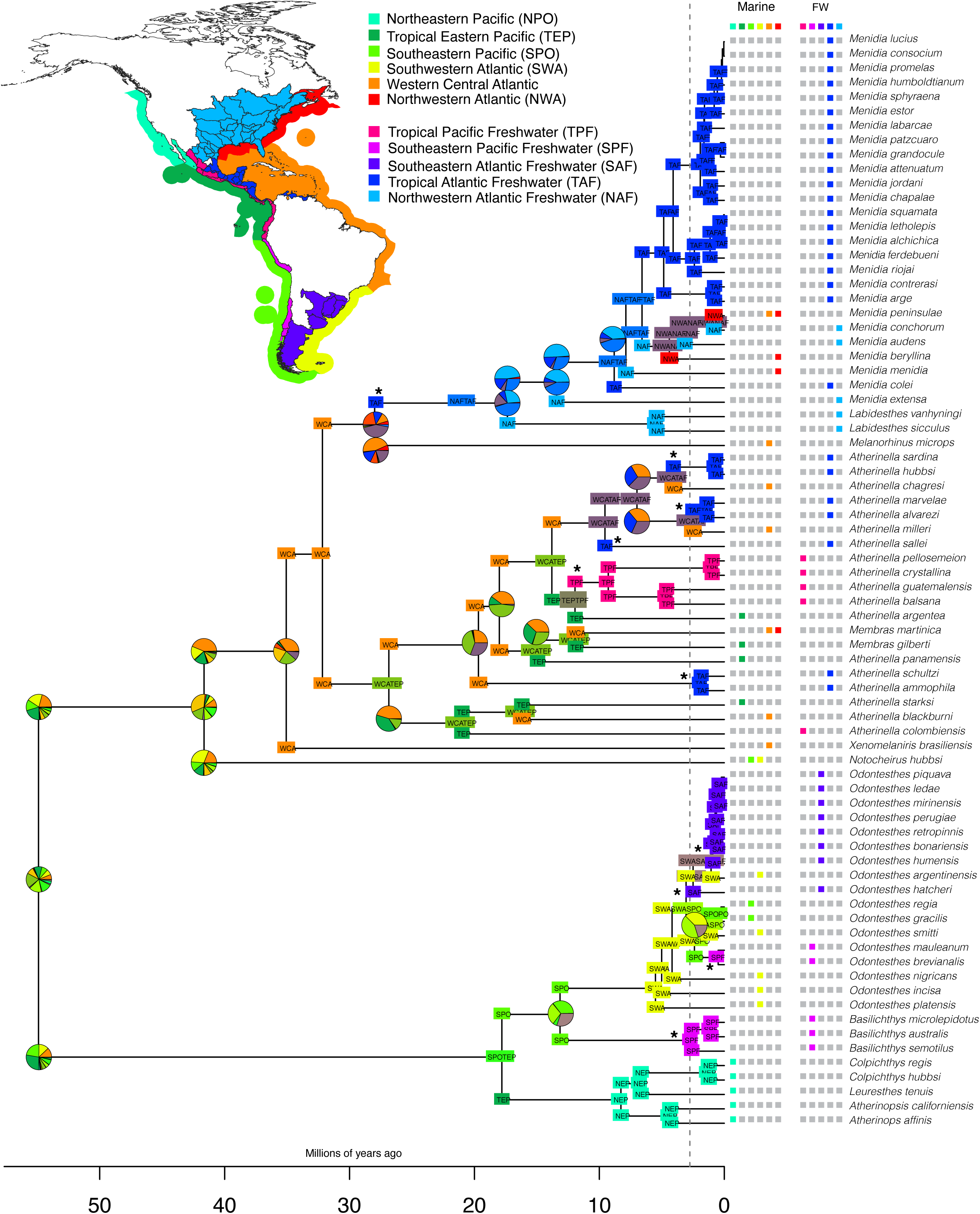
Biogeographic analysis of Atherinopsidae under the DEC model using the concatenated topology from Analysis 2 (Table 1). Dashed vertical line indicates the final closure of the Isthmus of Panama. Nodes where a single state made up >50% of the most likely area have only that area listed, but pie charts are shown for nodes where no single area accounted for the majority of likely states. Map of biogeographic regions is adapted from Marine- and Freshwater Ecoregions of the World areas (Abell et al., 2008; Spalding et al., 2007). Asterisks indicate instances of lineages becoming exclusively freshwater.

Central Atlantic and Tropical Eastern Pacific - *Membras gilberti and M. martinica, Atherinella blackburni* and *A. starksi* - diverged significantly earlier. The DEC model suggests just two reversals from freshwater habitats back into the marine environment in the marine species of *Menidia*.

We considered both the ML and MSC topologies of our analysis of Atherinoidei, as well as two different dispersal probabilities over the EPB (Figs. S24-S31). The ML topology puts Eastern Atlantic *Atherina* species as the sister group to all other atherinoids, and draws the ancestral area for the clade eastward toward the area that corresponded to the ancient Tethys Sea. This seaway connected the Mediterranean to the Indian Ocean throughout the Eocene. Ancestral nodes in the phylogeny are estimated to have been primarily in the Indian and Indo-Pacific Oceans (Fig. 6), though extant Indo-Pacific taxa are now concentrated in Atherinomoridae, with scattered representatives in other families. Alternatively, the MSC topology pulls out long-branched families Phallostethidae and Atherionidae as early diverging members of Atherinoidei, which shifts the center of origin westward toward the Indo-Pacific Ocean and Sunda shelf. The deeper nodes are primarily reconstructed in the Indo-Pacific Ocean, with less presence in the Western Indian Ocean.

**Fig. 6.**
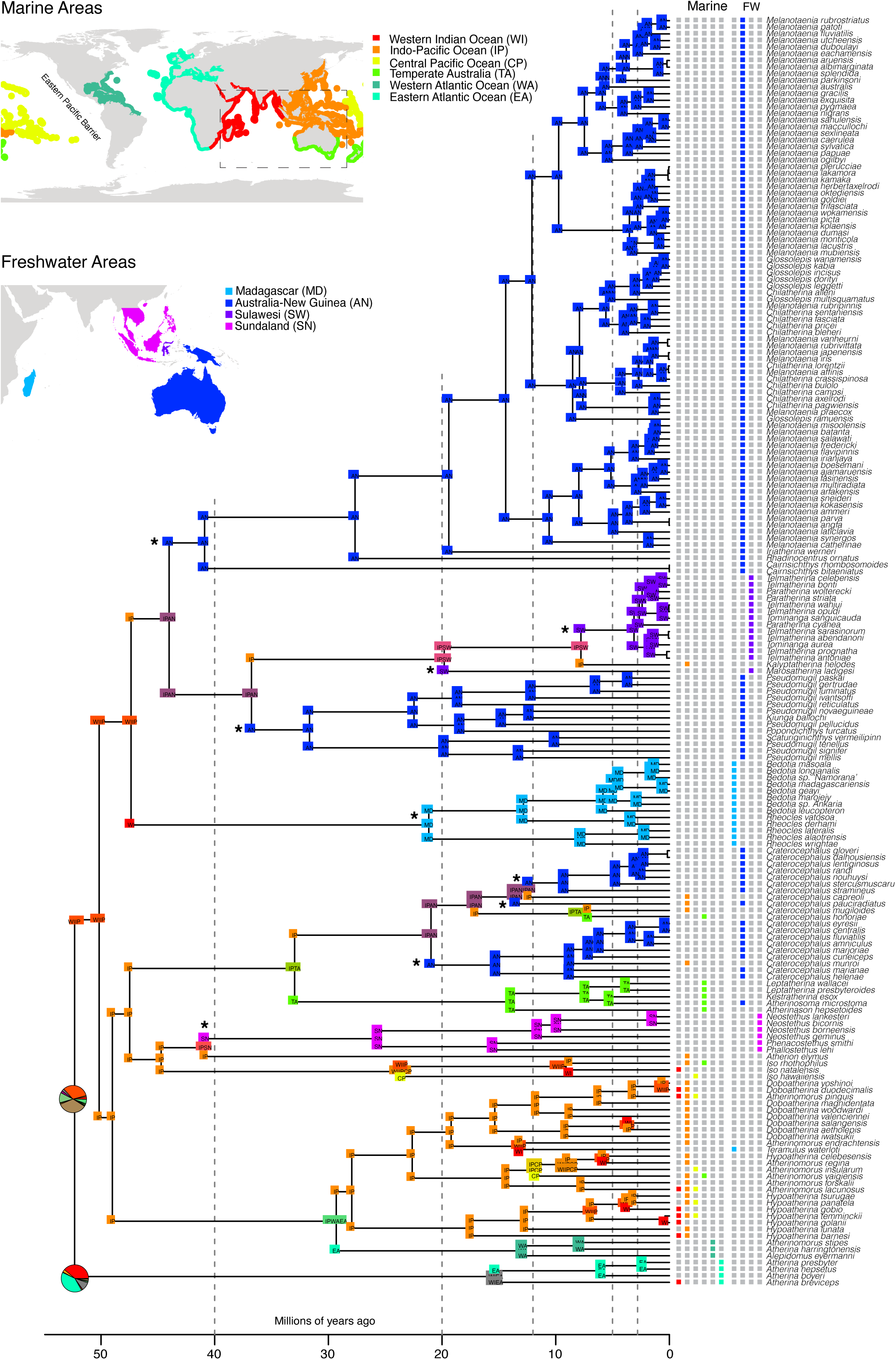
Biogeographic analysis of Atherinoidei under the DEC model using the concatenated topology from Analysis 2 (Table 1). Nodes where a single state made up >50% of the most likely area have only that area listed, but pie charts are shown for nodes where no single area accounted for the majority of likely states. This version of our model set a low dispersal probability across the EPB. Map of biogeographic regions is adapted from Marine- and Freshwater Ecoregions of the World areas (Abell et al., 2008; Spalding et al., 2007). Dashed vertical lines indicate from left to right: 1) the opening of the Makassar Strait ∼40 Ma, 2) the Sula Spur colliding with North Sulawesi ∼20 Ma, 3) the final closure of the Tethys Seaway ∼12 Ma, 4) the isolation of East Sulawesi from the Sahul Shelf ∼5 Ma, and 5) the final closure of the Isthmus of Panama ∼2.8 Ma. Asterisks indicate instances of lineages becoming exclusively freshwater.

The dispersal probability over the EPB affected the reconstruction of dispersal and extinction in the establishment of Atherinomoridae in the Western Atlantic - the only atherinoids present in the Americas, which are otherwise dominated by Atherinopisdae. When the EPB was considered a high barrier to dispersal, reconstructions favored ancestral dispersal, likely though the Tethys Seaway, from the Indo-Pacific to the Eastern Atlantic, then on to the Western Atlantic. When it was considered a low barrier to dispersal, the model favored the route through the Central Pacific.

## 4. Discussion

### 4.1 Limits of Phylogenomic Resolution in Atheriniformes

Ancient, rapid radiations in which branching times are short and do not allow for much accumulation of phylogenetic signal remain intractable in many groups despite the widespread application of phylogenomic methods (Arcila et al., 2021; Braun et al., 2019; Bravo et al., 2019). Atherinoidei exemplifies the challenges of these clades, with extremely short branches at the base of the tree. Across our 19 analyses – except for the unambiguous support for a clade containing Melanotaeniidae, Pseudomugilidae, and Telmatherinidae (MPT clade) – almost all families could be considered rogue taxa (Fig. 2). The position of Malagasy Rainbowfishes (Bedotiidae) is highly variable, sometimes forming the sister group to the MPT clade as other studies have supported (Sparks and Smith, 2004), but other times as the sister clade to Atherinimoridae (Fig. 2c,d) or Atherinidae (Fig. 2h). Long-branched families Atherionidae and Phallostethidae are alternatively deeply nested or early branching within Atherinoidei, depending on the analysis. A reduced dataset analyzed under the GHOST model, which accounts for heterotachy and should ameliorate long branch attraction artifacts, had high support for these families as nested in Atherinoidei. Excluding these long branches also did not affect the inferred ML topologies. Unfortunately, we are missing the monotypic Dentatherinidae, whose phylogenetic position based on morphology is debated (Ivantsoff et al., 1987; Parenti, 1984). Our attempt to get DNA from a formalin-fixed specimen following a protocol for natural history specimens was not successful (Ruane and Austin, 2017). The future inclusion of this taxon may help break some of these long branches, but is not likely to help in areas with low phylogenetic signals at the base of Atherinoidei.

Our support the need for a revised family-level classification of the group. Atherinidae *sensu* Dyer and Chernoff (1996), defined by three morphological characters, is not monophyletic in any of our analyses. The clades we obtain are similar to their subfamily classification with one exception: Craterocephalinae. While monophyly of Atherinidae was inferred in a molecular analysis by Campanella *et al*. (2013), the bootstrap support value they obtained was just 24 for this node. We unambiguously support three independent clades that can be recognized as families: 1) the family Atherinidae restricted to the genus *Atherina*, 2) Atherinomoridae, elevated from a subfamily supported by four morphological characters (Dyer and Chernoff, 1996); and 3) Craterocephalidae, whose circumscription differs from Craterocephalinae *sensu* Dyer & Chernoff by the inclusion of *Atherinason*, *Atherinasoma*, *Kestratherina*, and *Lepthatherina* (these genera group with *Atherina* in their morphological analysis). This new circumscription of Craterocephalidae is also more congruent with biogeography as all taxa are restricted to Australia, New Guinea, or their surrounding marine waters (Oyston et al., 2022). A revised family-level classification is provided at the end of this paper.

Resolution within Atherinopsidae using phylogenomic data was straightforward, and the relationships are consistent with those obtained with smaller mitochondrial or nuclear datasets (Bloom et al., 2012, 2009; Campanella et al., 2015), with some improvements on support values. Resolution for recent crown clades like *Odontesthes* is also robust, with the topology similar to that of a more detailed ddRADseq study using many more sites and individuals of that genus (Hughes et al., 2020). The position of *Melanorhinus microps* is among the few poorly supported nodes across all concatenated analyses (Figs. S1-S6), but always receives maximum support in MSC analyses (Figs. S14-S16).

### 4.2 Timing and Tempo of Atherinform Diversification

Atheriniformes have a poor fossil record, but many higher-level studies of fish phylogeny report remarkably consistent estimates for the age of origin for this group. These studies have put the age estimate in the latest Cretaceous between 71-76 Ma (Betancur-R. et al., 2013; Ghezelayagh et al., 2022; Hughes et al., 2018; Near et al., 2013). We used this as our soft maximum bound for our root calibration, and our median root age was 71.9 Ma (HPD: 66-77.5 Ma) regardless of whether we used the G75 ML topology or MSC topology as a constraint.

Atherinopsidae and Atherinoidei almost simultaneously next split at a median age of 54-52 Ma, the period directly following the Paleocene-Eocene Thermal Maximum at ∼55.8 Ma, an extreme warming event that caused marine mass extinctions in some groups of fishes (Arcila and Tyler, 2017). Atherinoidei radiated rapidly during the Eocene climatic optimum, the warmest period of the Cenozoic when there was little ice at the poles. The relatively high sea level at this time created many shallow seaways that may have facilitated rapid dispersal by this group whose marine members favor coastal environments.

### 4.3 Biogeographic Analysis Reveals Erased Marine Ancestry

Atheriniformes are exceptional at transitioning between marine and freshwater biomes. Studies often code these as binary states to understand habitat transitions in Atheriniformes or across the Tree of Life of fishes (Betancur-R et al., 2015; Bloom et al., 2013; Campanella et al., 2015; Rabosky, 2020), however this erases the spatial context of these ecological transitions.

Atheriniformes diversified after the breakup of ancient supercontinents, thus their global distribution can only be explained by extensive marine dispersal and colonization of freshwater habitats. We estimated ancestral areas under the DEC model with some biologically realistic limitations, and since even marine atheriniform species tend to have limited regional distributions they are well suited to a formal biogeographic analysis. By limiting dispersal so it could only occur between adjacent areas and limiting the maximum number of areas a species could inhabit to the largest number of areas observed in extant species we were able to code the large number of marine and freshwater ecoregions that atheriniform taxa inhabit. Broadly, these analyses suggest widespread marine ancestry erased by extinction, greatly increasing the number of likely marine-to-freshwater transitions when compared to an analysis of binary states (Campanella et al., 2015).

Atherinopsidae is widespread throughout the Americas (Fig. 5), dominated by the anti-tropical Atherinopsinae and the primarily tropical Menidiinae. This has long been attributed to middle-Miocene warming (White, 1986), and our divergence time estimates are generally congruent with that hypothesis. Our biogeographical reconstruction suggests that the ancestral Atherinopsinae node was present in the Tropical Eastern Pacific, where no species currently inhabit. Comparable TEP absences occur in other broadly distributed marine fishes, including the bluntnose lizardfish (*Trachinocephalus*), the cobia (*Rachycentron*), and the longfin escolar (*Scombrolabrax*), all of which are present in the Western Atlantic and therefore likely occupied the Eastern Pacific prior to the final closure of the Isthmus of Panama. Their continued absence suggests that regional extinction, compounded by the Eastern Pacific Barrier preventing recolonization, can obscure ancestral distributions. Deeper nodes in Atherinopsidae have more ambiguous biogeographic reconstructions, and are also likely obscured by extinction.

Phylogenetic uncertainty among atherinoid lineages makes the geographic origin of this clade more difficult to infer. The presence of two extinct atherinoid families Mesogasteridae and Rhamphognathidae from the Eocene (Ypresian) Monte Bolca lagerstätten suggests an ancient presence in the Tethys Sea, though their relationship to extant families is unknown (Bannikov, 2008). Our topologies that place *Atherina* as the sister to all other atherinoids pulls the origin of this clade eastward, toward what would have been the Tethys Sea (Fig. 6). However, our analysis of the MSC topology favors a more western origin in the Indo-Pacific-Sundaland area (Figs. S24-S27). Regardless of phylogenetic and fossil record uncertainty, formal biogeographic analysis considering marine regions and freshwater areas better reflects the marine-to-freshwater transitions in this group than a simple two-state coding can capture. This is evident in the strongly supported clade of Melanotaeniidae, Telmatherinidae, and Pseudomugilidae, comprised almost exclusively of freshwater taxa. Melanotaeniidae and Pseudomugilidae primarily inhabit freshwater habitats of the Sahul shelf, now Australia and New Guinea. No extant melanotaeniids inhabit marine waters, however a few pseudomugilids (e.g. *Pseudomugil halophilus* and *Pseudomugil inconspicuus*) do inhabit marine and brackish habitat. Unfortunately, those taxa are missing from our analysis. Telmatherinidae is endemic to the Wallacean island of Sulawesi, a composite island formed of both the Asian and Australian plates, except for marine *Kalyptatherina helodes*. The DEC model with biogeographical coding and limiting dispersal to adjacent areas reconstructed at least three separate transitions from the Indo-Pacific Ocean to establish these families in freshwater habitats (Fig. 6). In a binary marine-freshwater coding scheme, the ancestor of these three families is always inferred to be of freshwater origin (Campanella et al., 2015).

Three species of Atherinomoridae inhabit the Western Atlantic, the only atherinoids present in the Americas. This lineage is sister to a predominantly Indo-Pacific clade. Whether the biogeographic model favors dispersal and extinction of a marine ancestor through an East Atlantic/Tethyan route or through the Central Pacific depends largely on how we consider the EPB as a barrier to dispersal. Models where the EPB was considered a high barrier to dispersal favored the former reconstruction (Fig. 6), while models where the EPB was considered a low barrier to dispersal favored the latter (Figs. S24-31). Notably, these three Atherinomoridae species diversified after the final closure of the Tethys seaway.

### 4.4 Revised Family-Level Classification of Atheriniformes

Order **Atheriniformes**

Suborder **Atherinopsoidei**

Family **Atherinopsidae** Fitzinger 1873 Type genus: *Atherinopsis* Girard 1854

Subfamily **Atherinopsinae** Fitzinger 1873 *Atherinops, Atherinopsis, Basilichthys, Colpichthys, Leuresthes, Odontesthes*

Subfamily **Notocheirinae** Shultz 1950 *Notocheirus*

Subfamily **Menidiinae** Schultz 1948 *Atherinella, Labidesthes, Melanorhinus, Membras, Menidia, Xenomelaniris*

Suborder **Atherinoidei**

Family **Atherinidae** Risso 1827 Type genus: *Atherina* Linnaeus 1758 *Atherina*

Family **Atherionidae** Schultz 1948 Type genus: *Atherion* Jordan & Starks 1901 *Atherion*

Family **Isonidae** Rosen 1964 Type genus: *Iso* Jordan & Starks 1901 *Iso*

Family **Phallostethidae** Regan 1916 Type genus: *Phallostethus* Regan 1913

*Gulaphallus*\**, Neostethus, Phallostethus, Phenacostethus*

Family **Atherinomoridae** Dyer & Chernoff 1996 (**New Circumscription) Type genus: *Atherinomorus* Fowler 1903 *Alepidomus, Atherinomorus, Doboatherina, Hypoatherina, Teramulus*

Family **Craterocephalidae** Dyer & Chernoff 1996 *New Circumscription Type genus: *Craterocephalus* McCulloch 1912 *Atherinason, Atherinasoma, Craterocephalus, Kestratherina, Leptatherina, Sashatherina*\*

Family **Dentatherinidae** Patten & Ivantsoff 1983 Type genus: *Dentatherina* Patten & Ivantsoff 1983 *Dentatherina*\*

Family **Bedotiidae** Jordan & Hubbs 1919 Type genus: *Bedotia* Regan 1903 *Bedotia, Rheocles*

Family **Pseudomugilidae** Kner 1867 Type genus: *Pseudomugil* Kner 1866 *Kiunga, Pseudomugil, Scaturiginichthys*

Family **Telmatherinidae** Munro 1958 Type genus: *Telmatherina* Boulenger 1897

*Kalyptatherina, Marosatherina, Paratherina, Telmatherina, Tominanga*

Family **Melanotaeniidae** Gill 1894 Type genus: *Melanotaenia* Gill 1862 *Chilatherina, Glossolepis, Melanotaenia, Cairnsichthys, Rhadinocentrus, Iriatherina, Pelangia*

*Incertae sedis*: *Bleheratherina* Aarn & Ivansoff 2009

## Acknowledgements

We would like to thank the many researchers, curators and collections managers who provided us with tissues for this project, including: Carole Baldwin, Lynne Parenti, and Diane Pitassy (USNM); the Smithsonian Tropical Research Institute (STRI); Mariano Castro-González (UNMPD); Luis Malabarba (UFRGS); Mark Sabaj (ANSP); Prosanta Chakrabarty (LSUMZ); Will White and John Pogonowski (CSIRO); Nathan Lovejoy (University of Toronto); Masaki Miya (Natural History Museum, Chiba); David Reznick (UC Riverside). Analyses were conducted on the NC State University high performance computing cluster Hazel. This work was supported by the following National Science Foundation grants: DEB-1754627 (to DDB), DEB-1932759 (to RBR), and DEB 1541554 (to GO).

## Data Availability

Sequence data is available at NCBI Bioproject number PRJXXXXX (submission pending). Supplemental figures, tables, sequence alignments, tree files, and R Code are available on Zenodo doi:10.5281/zenodo.19255610.

