## Supplemental Figures for "Phylogenomics, Biogeography, and a New Family-level Classification of Silversides, Rainbowfishes, and Allies (Teleostei: Atheriniformes)"

**Fig. S1: Maximum likelihood phylogeny of all nucleotides (Table 1: analysis 1). Node values indicate ultrafast boostrap support.**

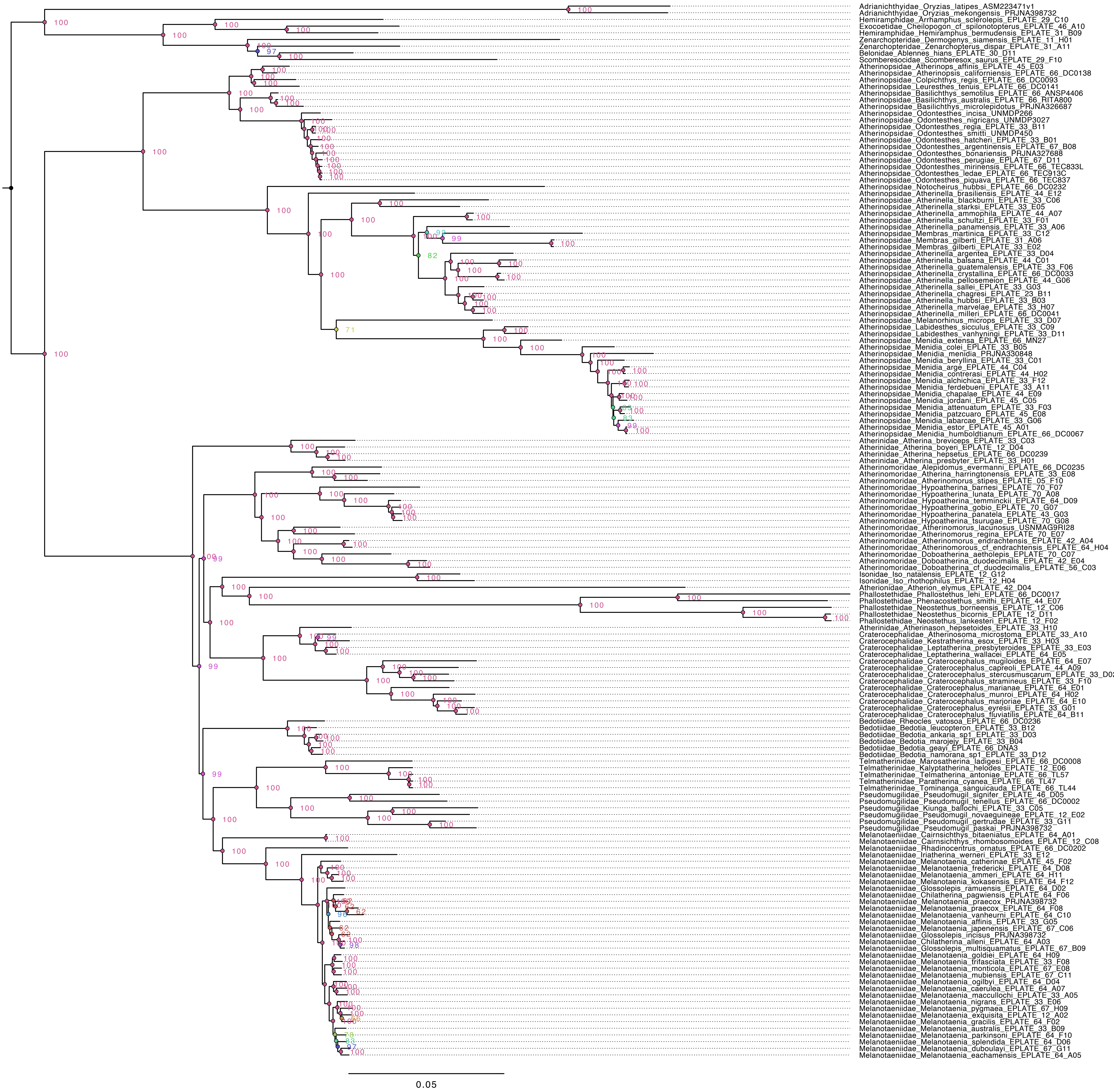

**Fig. S2: Maximum likelihood phylogeny of G75 nucleotide matrix (Table 1: analysis 2). Node values indicate ultrafast boostrap support.**

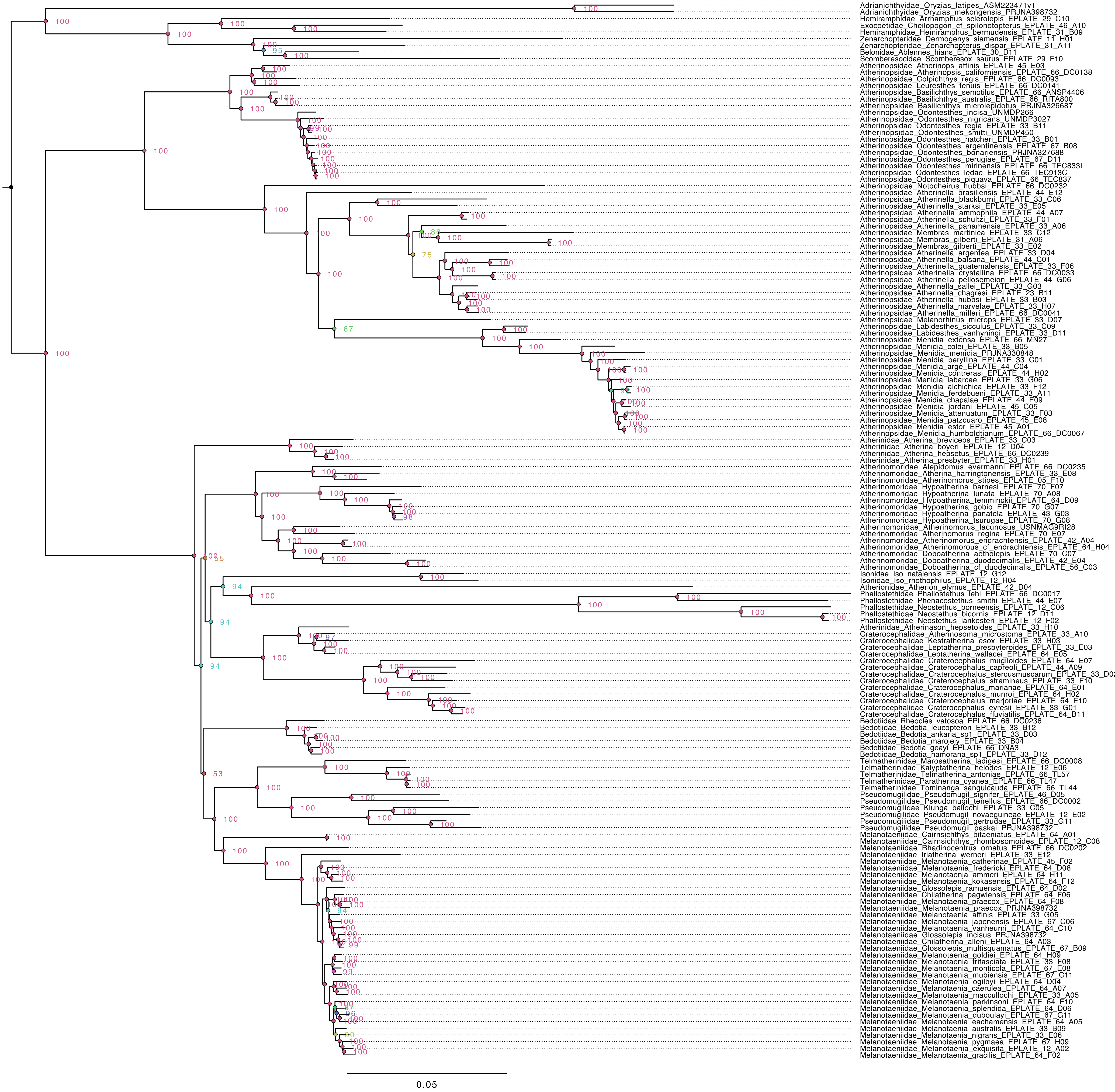

**Fig. S3: Maximum likelihood phylogeny of G90 nucleotide matrix (Table 1: analysis 3). Node values indicate ultrafast boostrap support.**

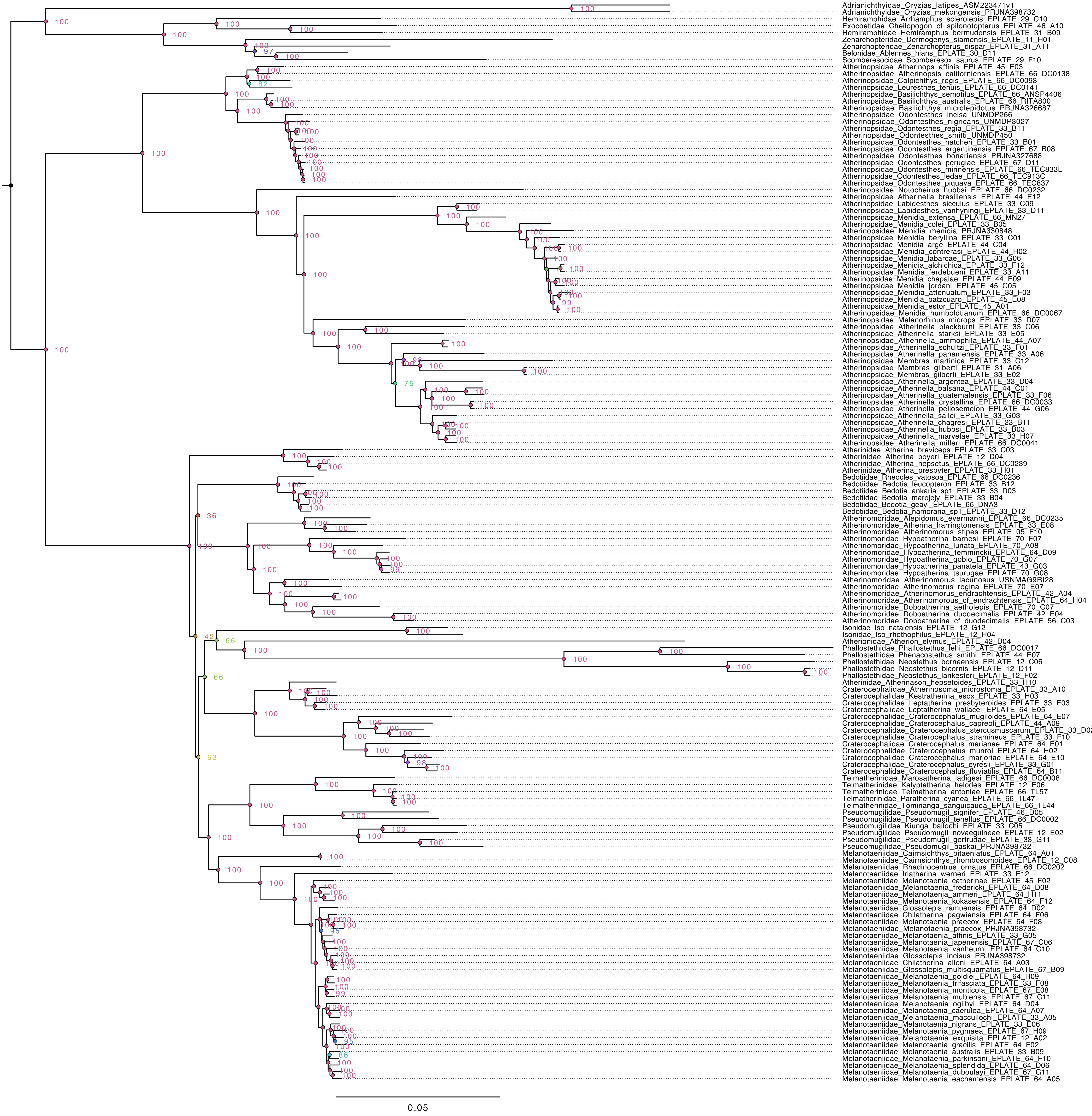

**Fig. S4: Maximum likelihood phylogeny of all amino acids (Table 1: analysis 4). Node values indicate ultrafast boostrap support.**

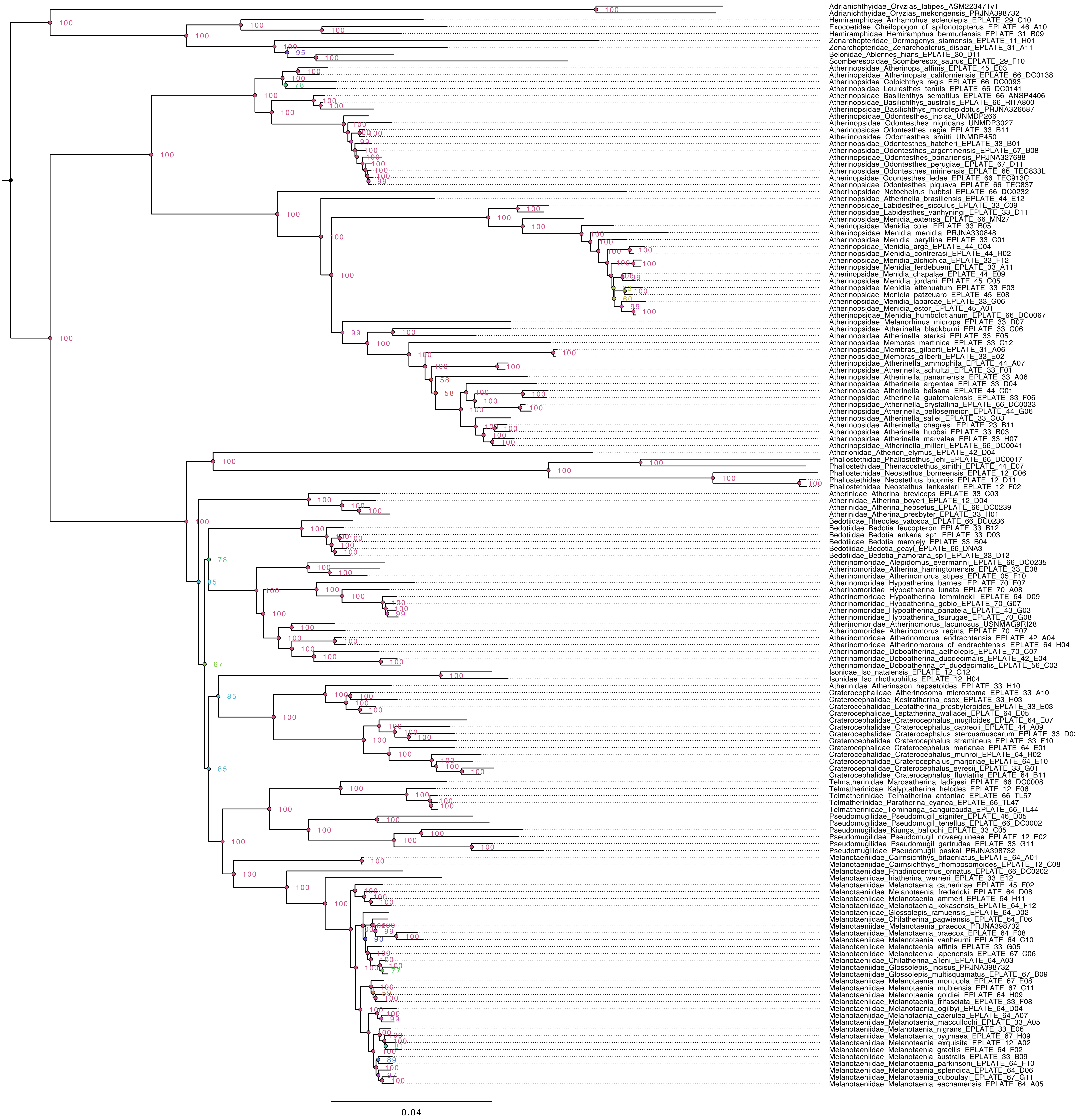

**Fig. S5: Maximum likelihood phylogeny of G75 amino acid matrix (Table 1: analysis 5). Node values indicate ultrafast boostrap support.**

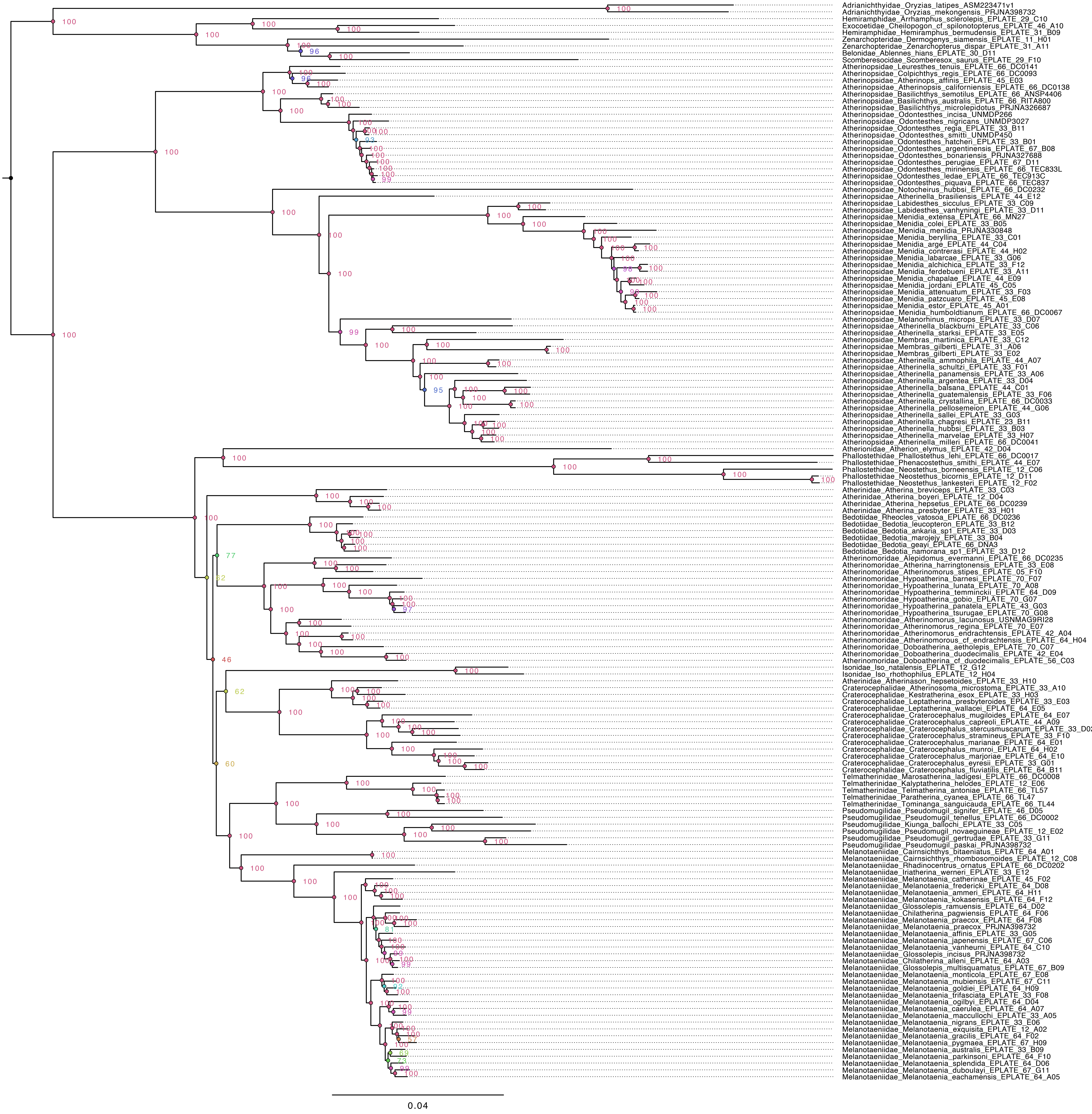

**Fig. S6: Maximum likelihood phylogeny of G90 amino acid matrix (Table 1: analysis 6). Node values indicate ultrafast boostrap support.**

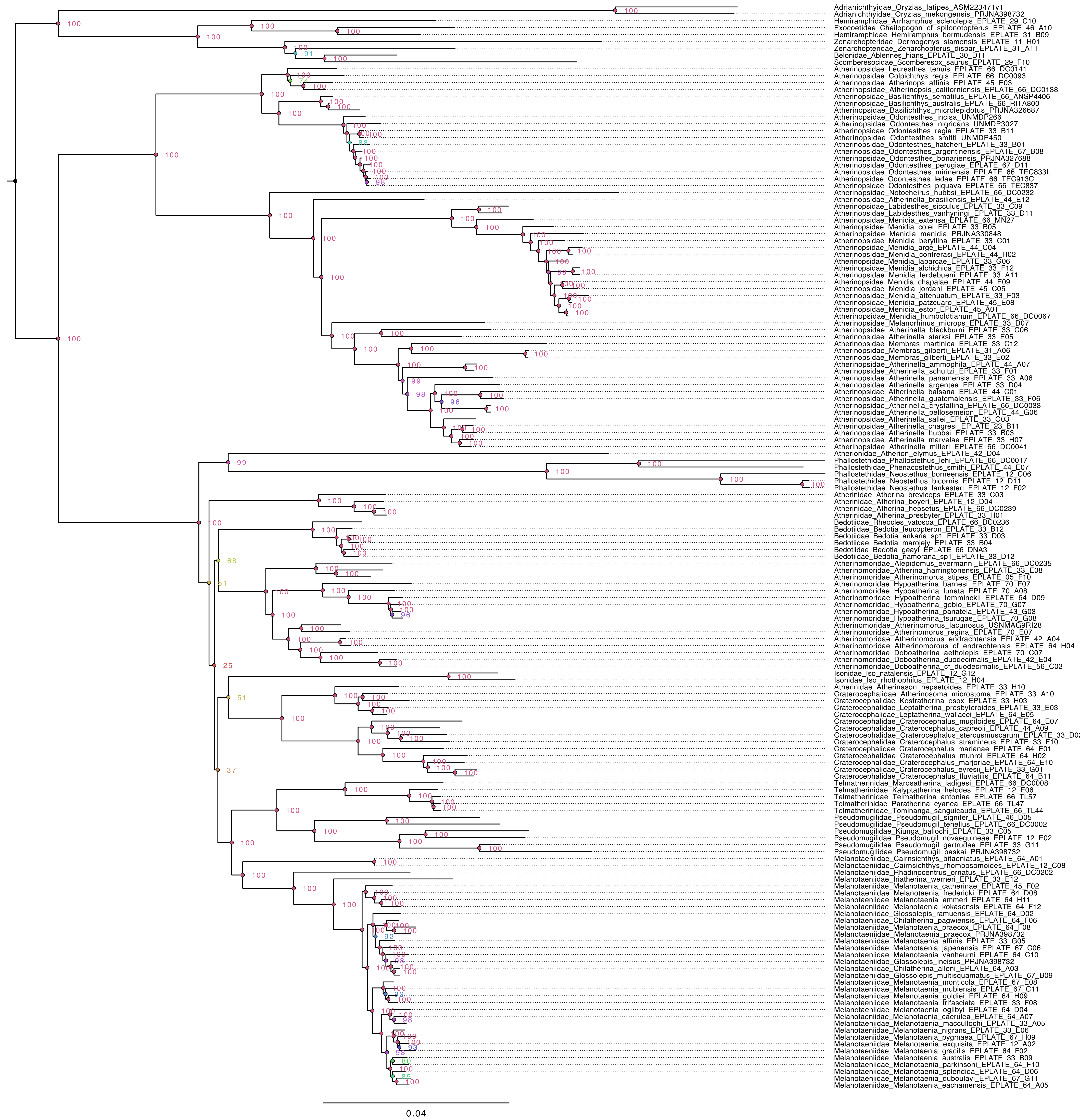

**Fig. S7: Maximum likelihood phylogeny of Atherinoidei using all nucleotides with Atherionidae excluded and reduced outgroups (Table 1: analysis 7). Node values indicate ultrafast boostrap support.**

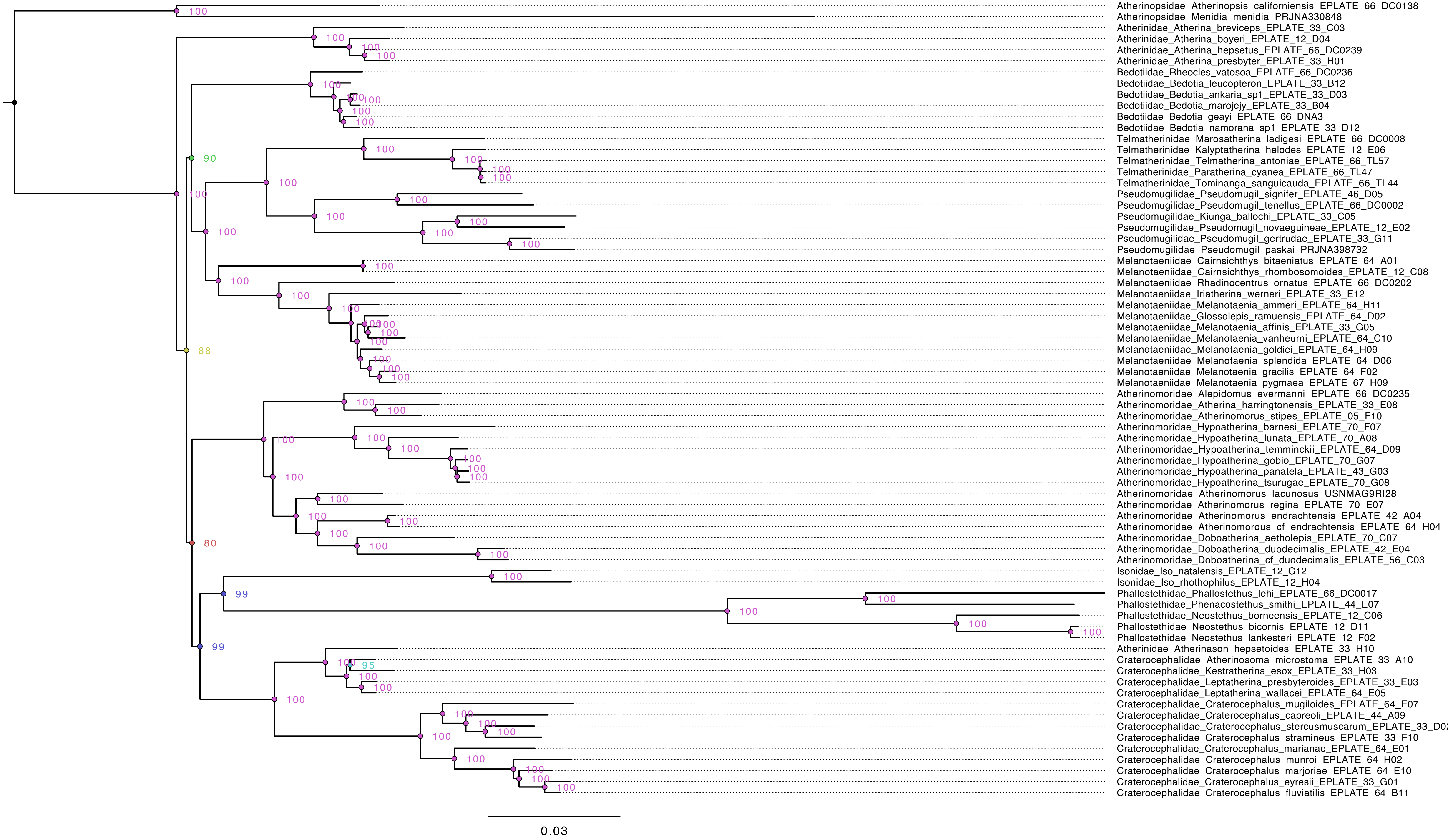

**Fig. S8: Maximum likelihood phylogeny of Atherinoidei using all nucleotides with Isonidae excluded and reduced outgroups (Table 1: analysis 8). Node values indicate ultrafast boostrap support.**

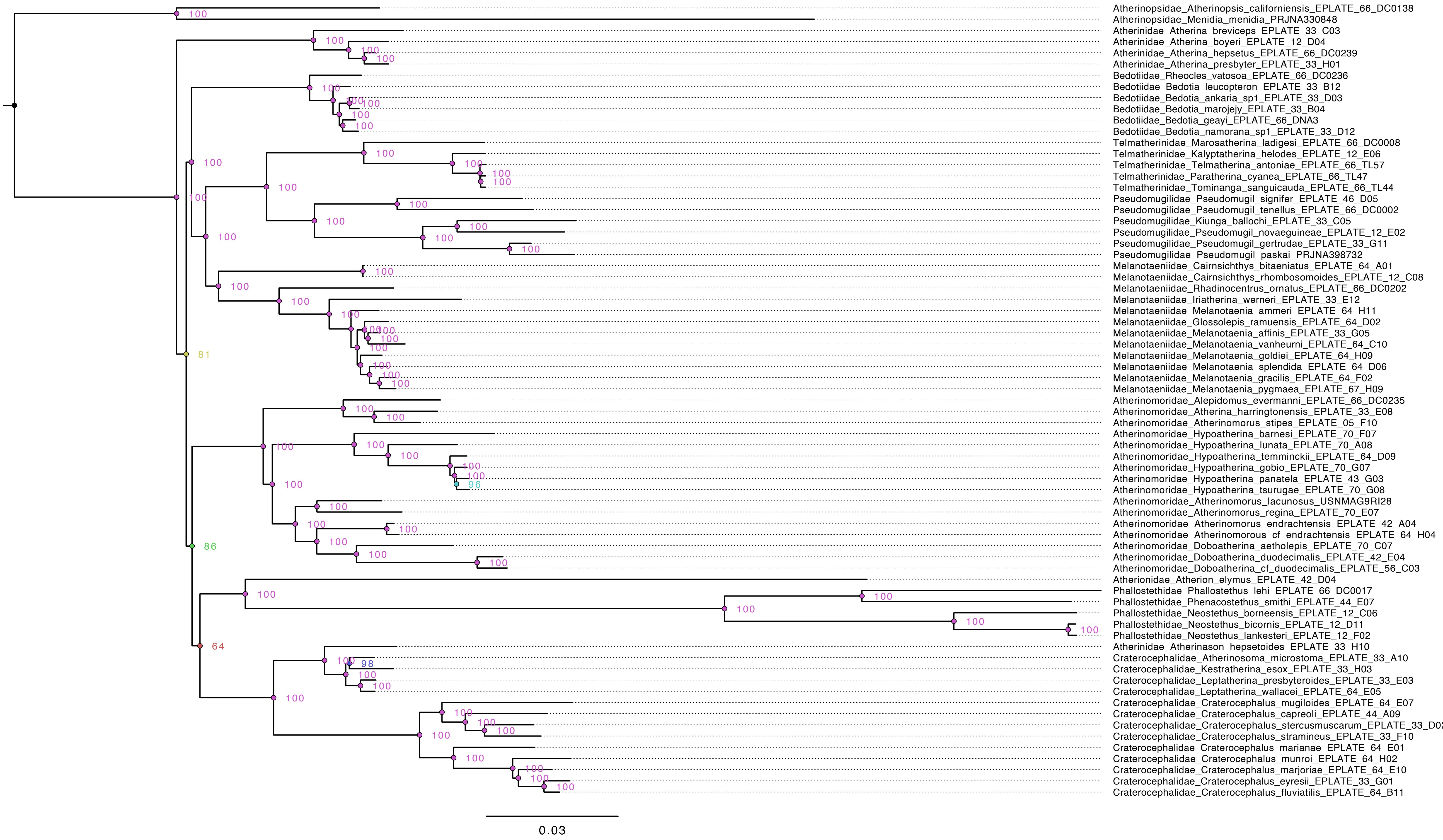

**Fig. S9: Maximum likelihood phylogeny of Atherinoidei using all nucleotides with Phallostethidae excluded and reduced outgroups (Table 1: analysis 9). Node values indicate ultrafast boostrap support.**

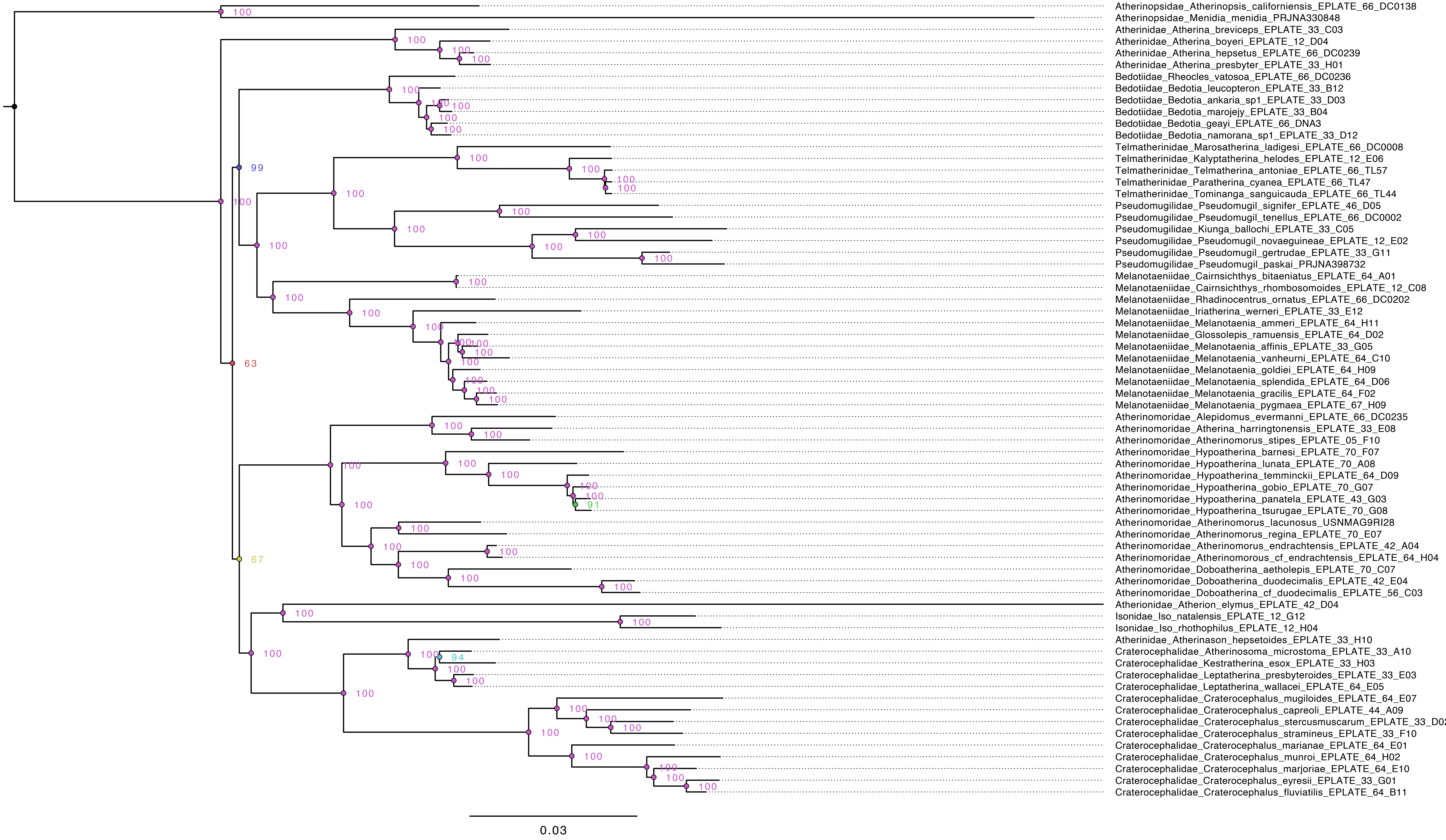

**Fig. S10: Maximum likelihood phylogeny of Atherinoidei using all nucleotides with reduced outgroups (Table 1: analysis 10). Node values indicate ultrafast boostrap support.**

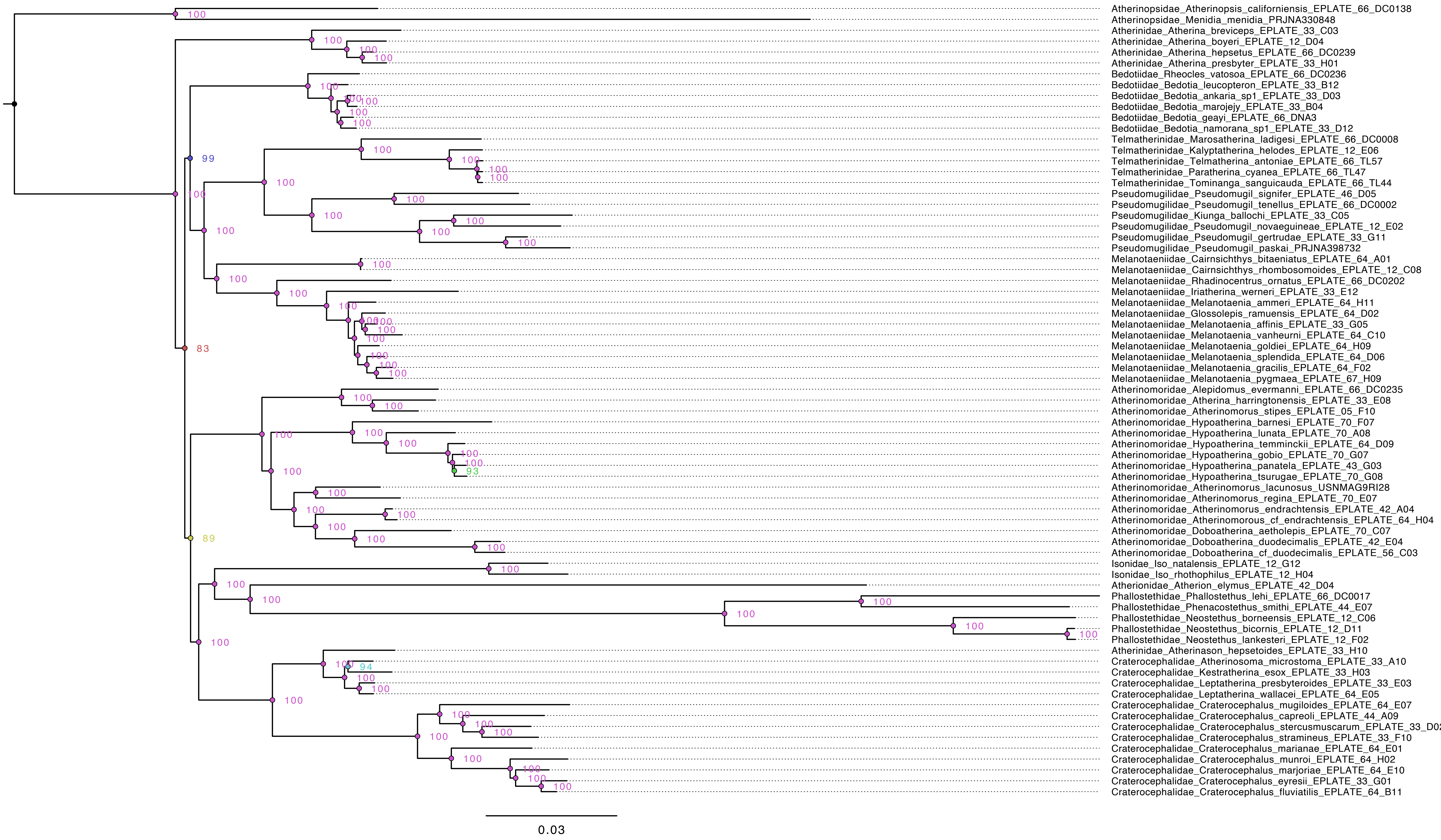

**Fig. S11: Maximum likelihood phylogeny of Atherinoidei using the G75 matrix with reduced outgroups (Table 1: analysis 11). Node values indicate ultrafast boostrap support.**

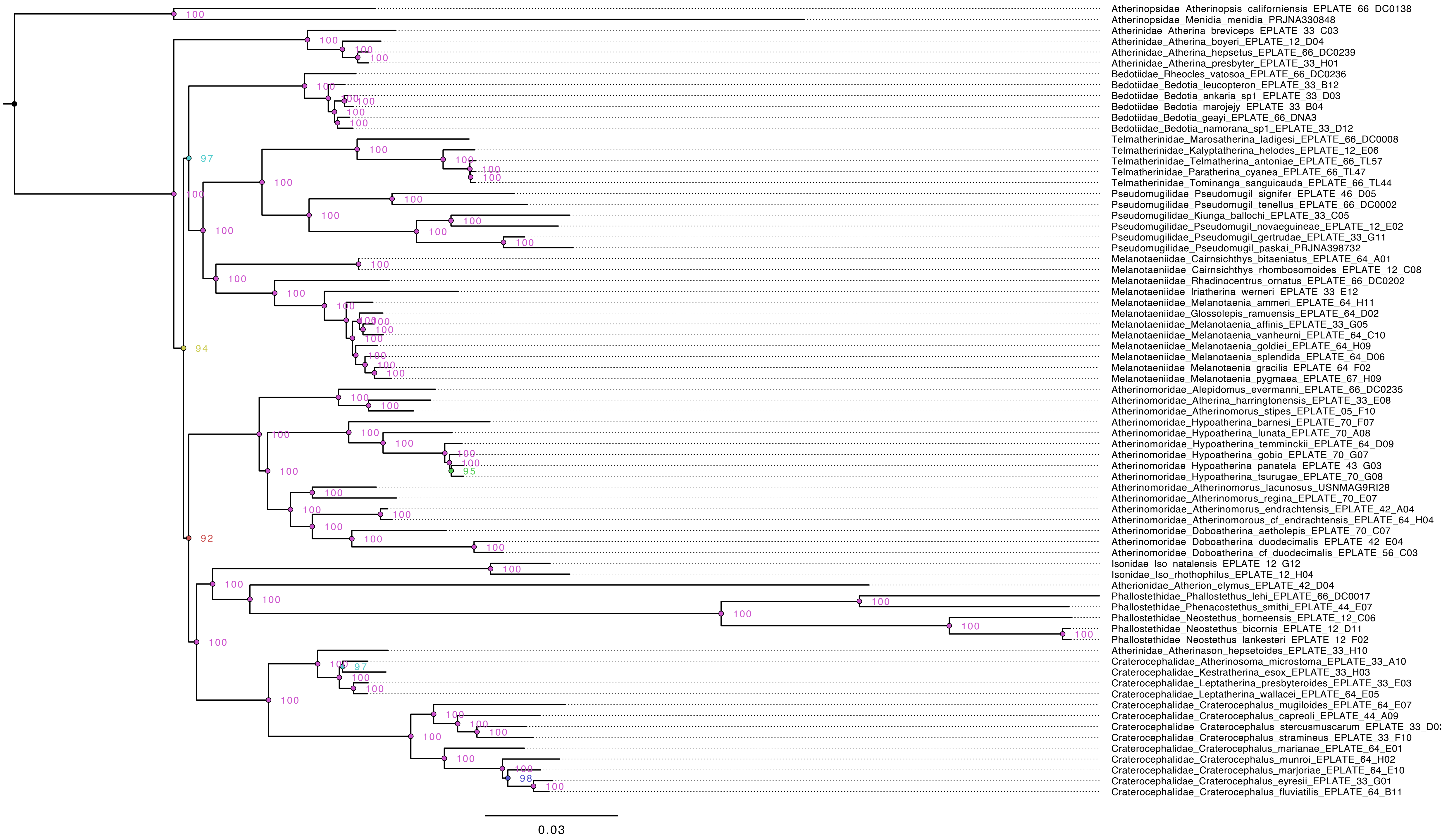

**Fig. S12: Maximum likelihood phylogeny of Atherinoidei using the G90 matrix with reduced outgroups (Table 1: analysis 12). Node values indicate ultrafast boostrap support.**

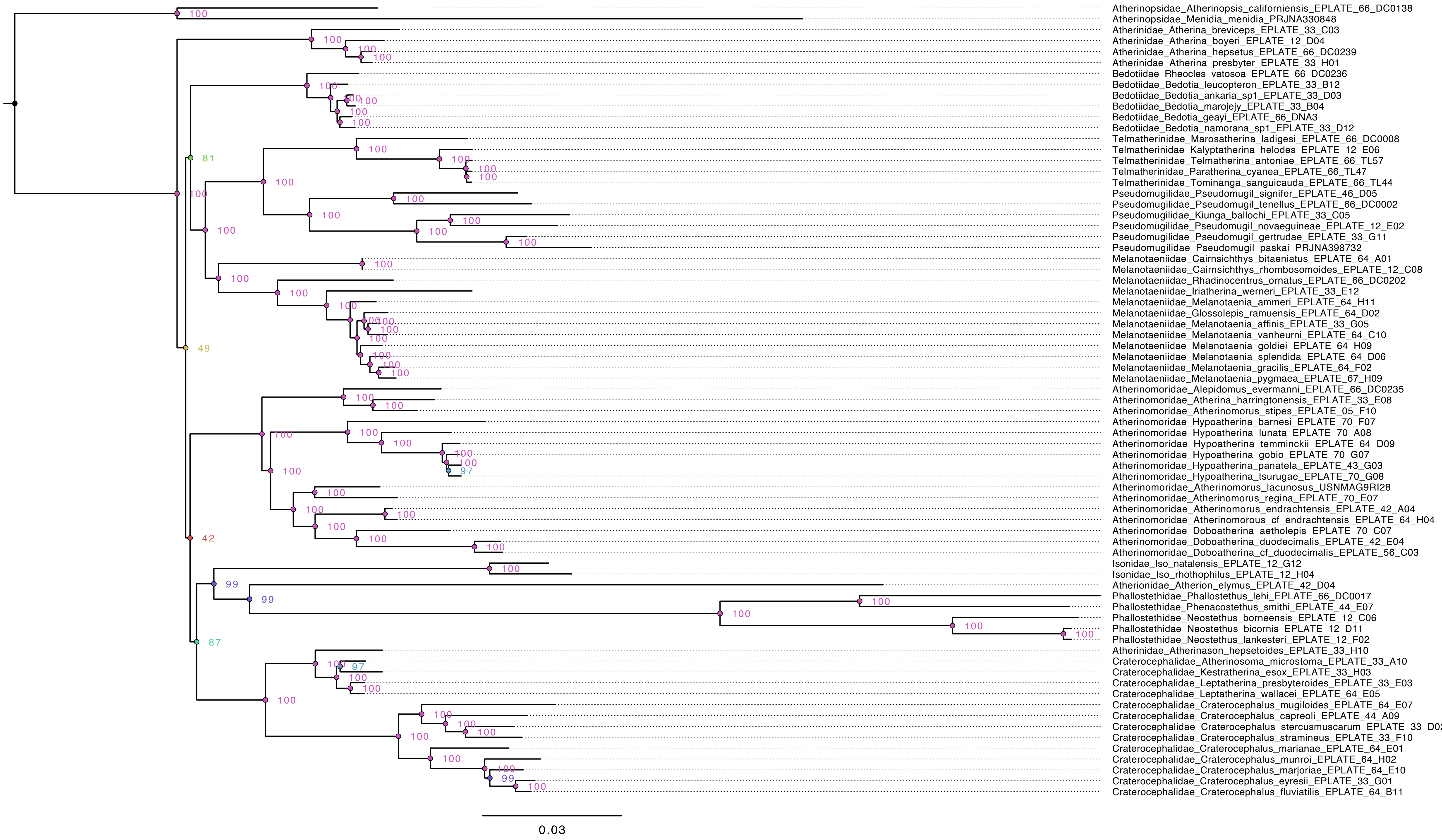

**Fig. S13: Maximum likelihood phylogeny of Atherinoidei using the G75 matrix with reduced outgroups under the GHOST model (Table 1: analysis 13). Node values indicate ultrafast boostrap support.**

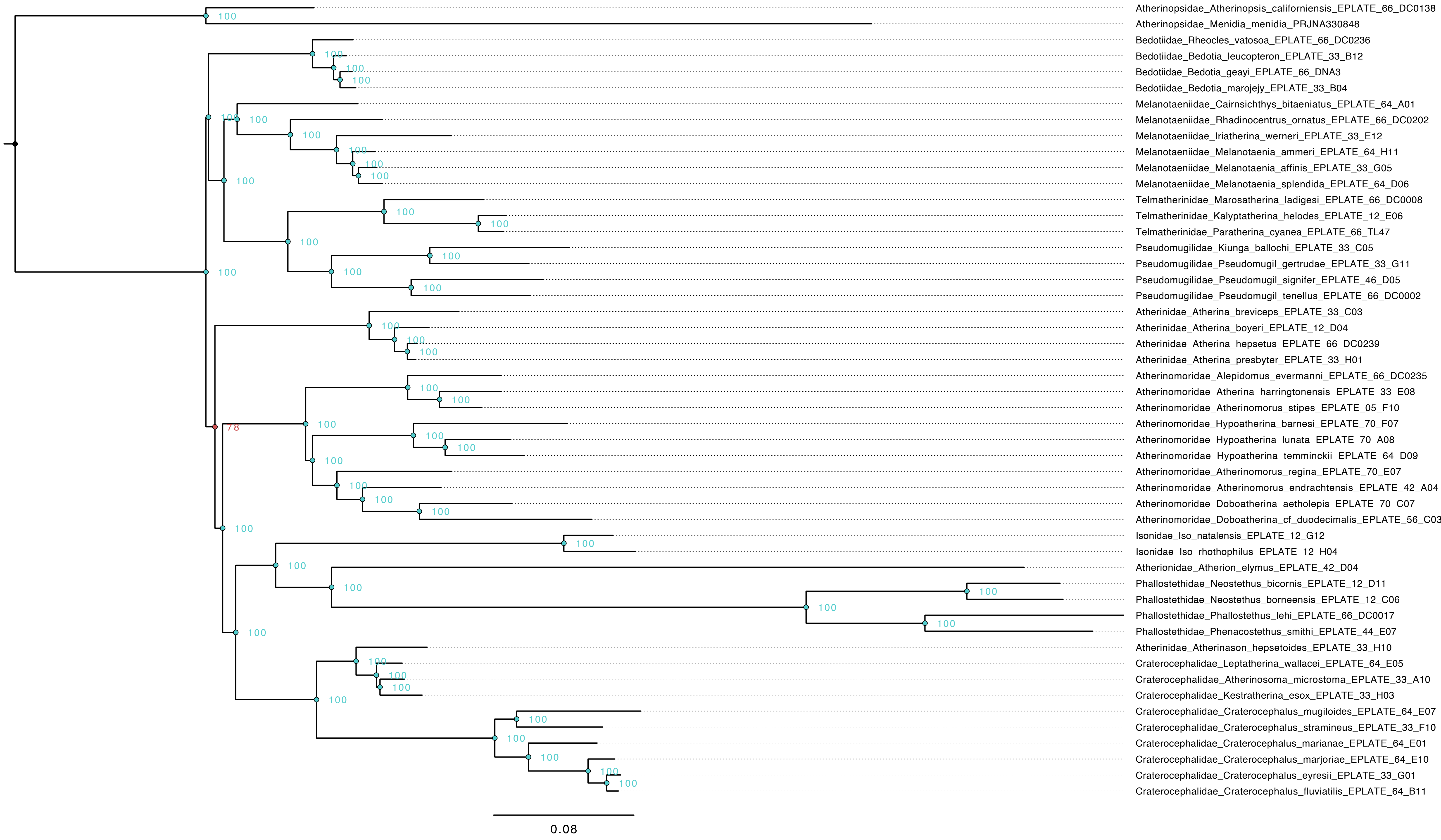

**Fig. S14: ASTRAL species tree estimated from all gene trees (Table 1: analysis 14). Node values indicate local posterior probability.**

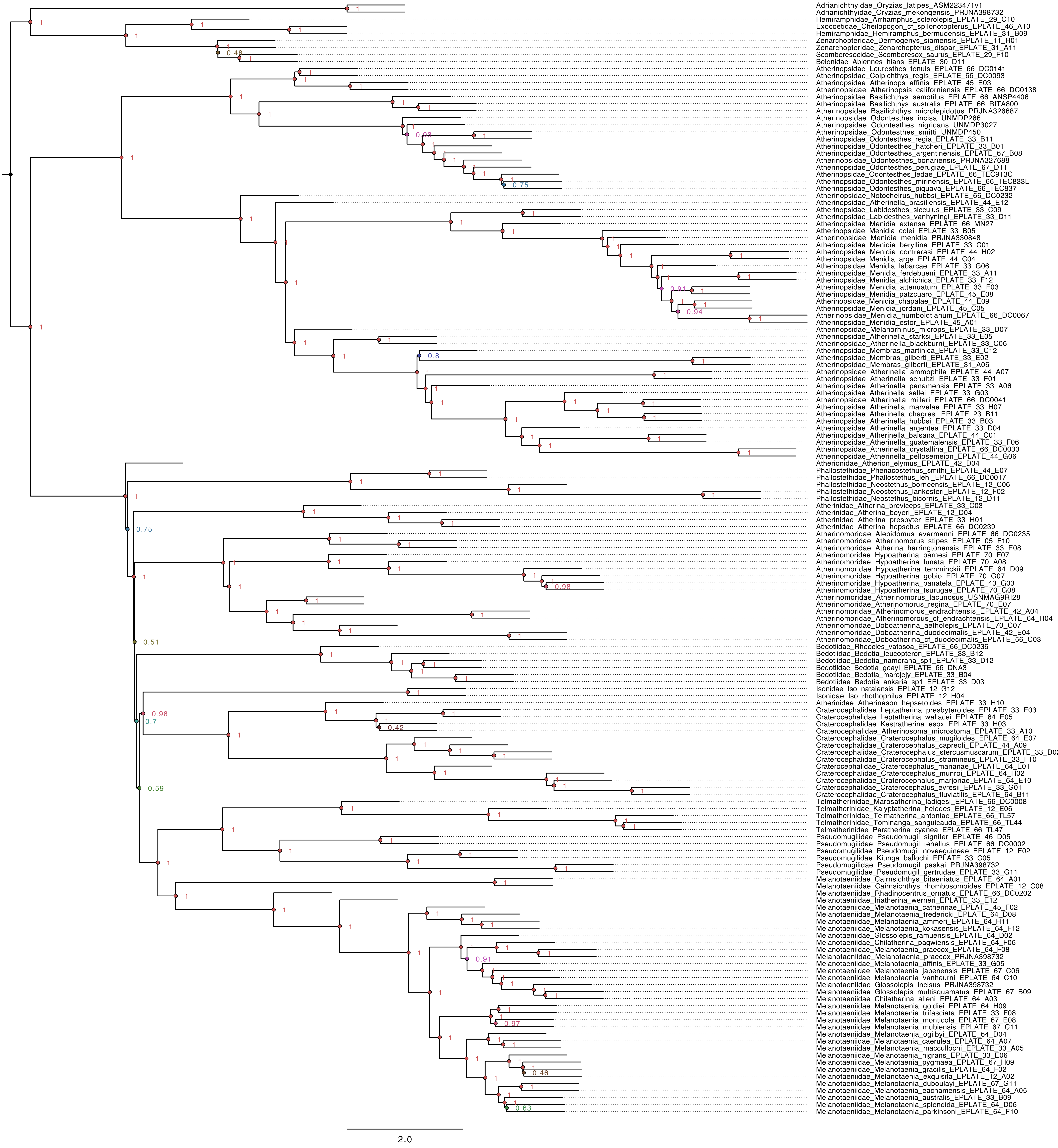

**Fig. S15: ASTRAL species tree estimated from G75 gene trees (Table 1: analysis 15). Node values indicate local posterior probability.**

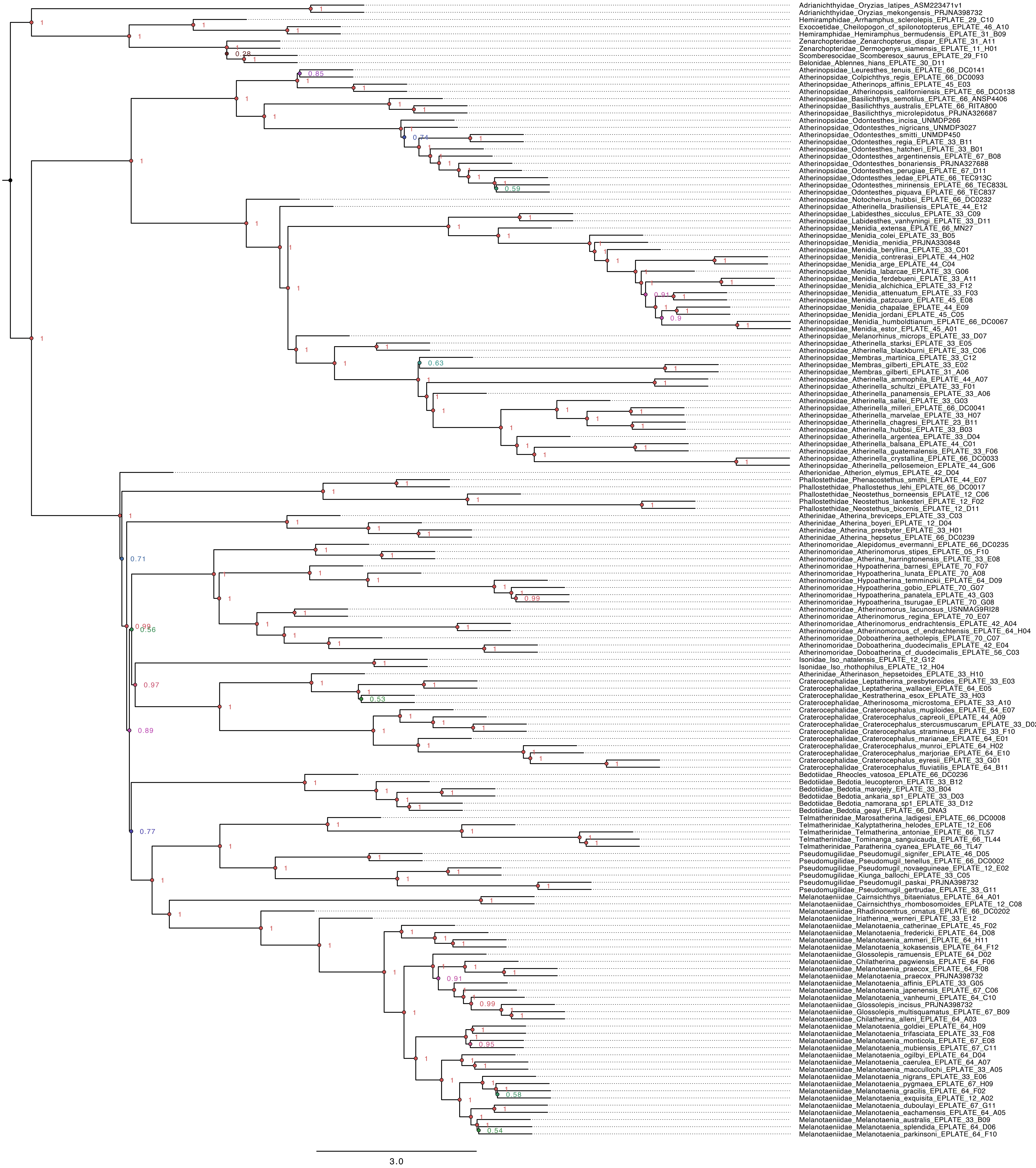

**Fig. S16: ASTRAL species tree estimated from G90 gene trees (Table 1: analysis 16). Node values indicate local posterior probability.**

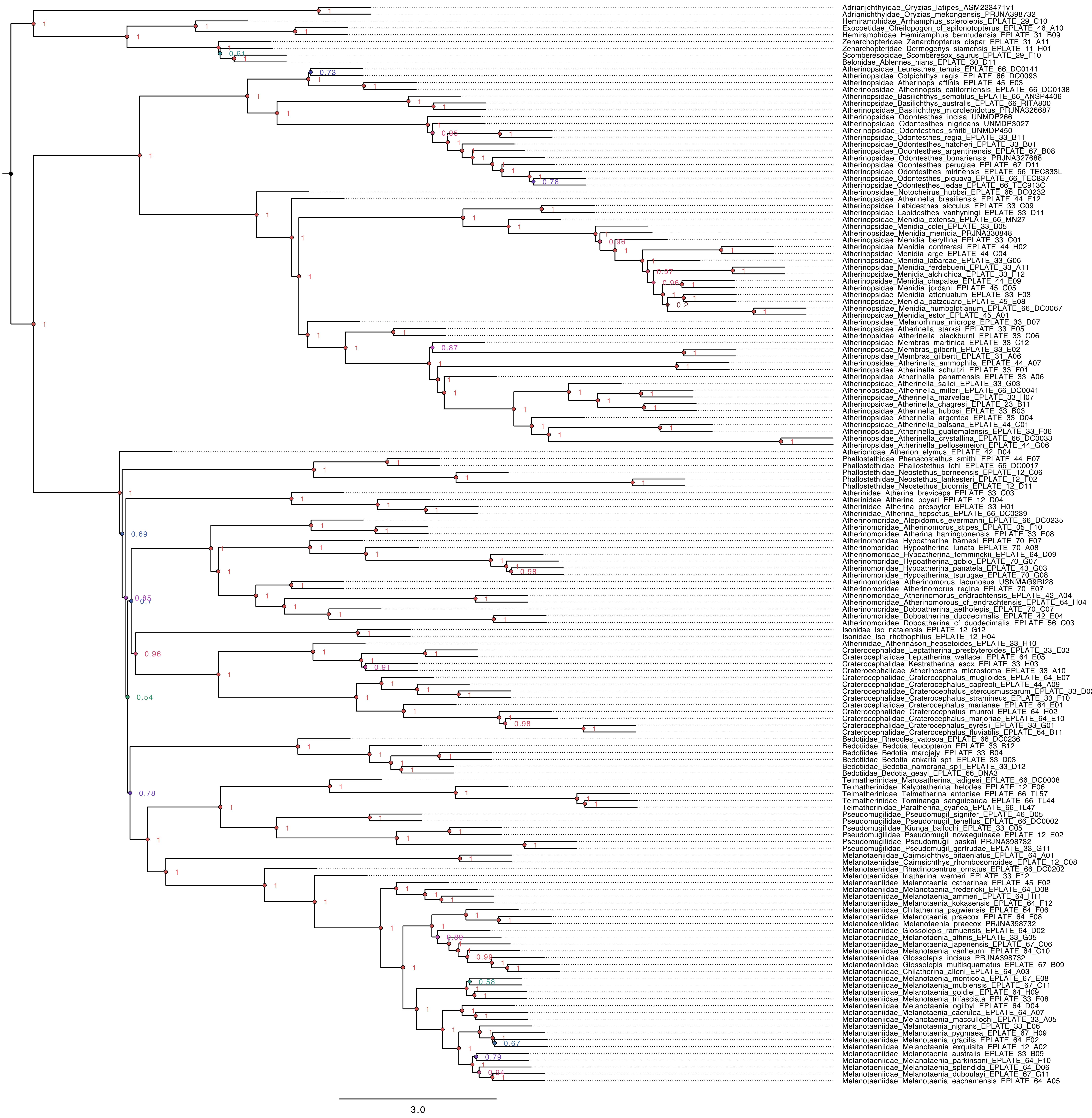

**Fig. S17: ASTRAL species tree of Atherinoidei estimated from all gene trees with reduced outgroups (Table 1: analysis 17). Node values indicate local posterior probability.**

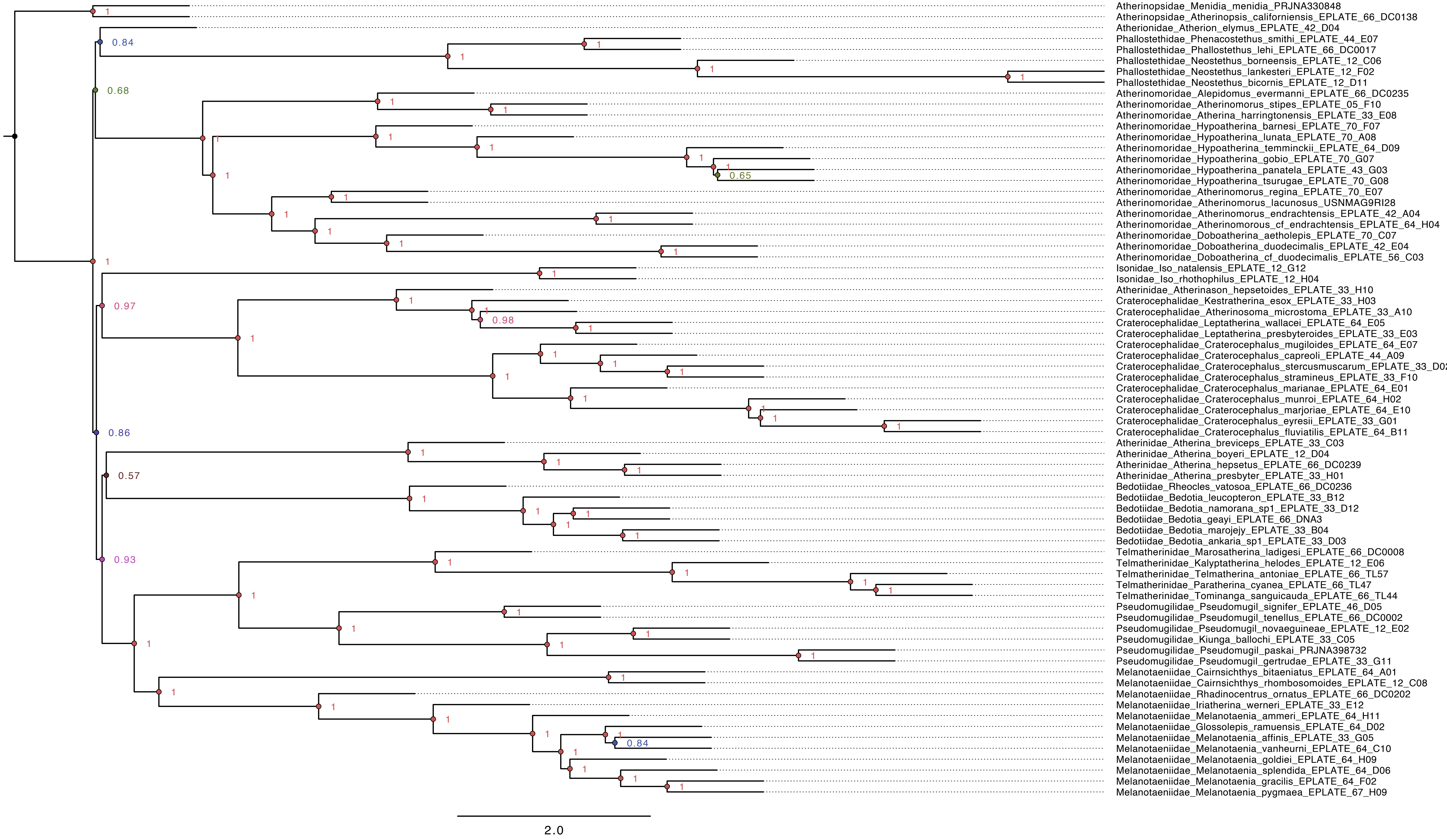

**Fig. S18: ASTRAL species tree of Atherinoidei estimated from G75 gene trees with reduced outgroups (Table 1: analysis 18). Node values indicate local posterior probability.**

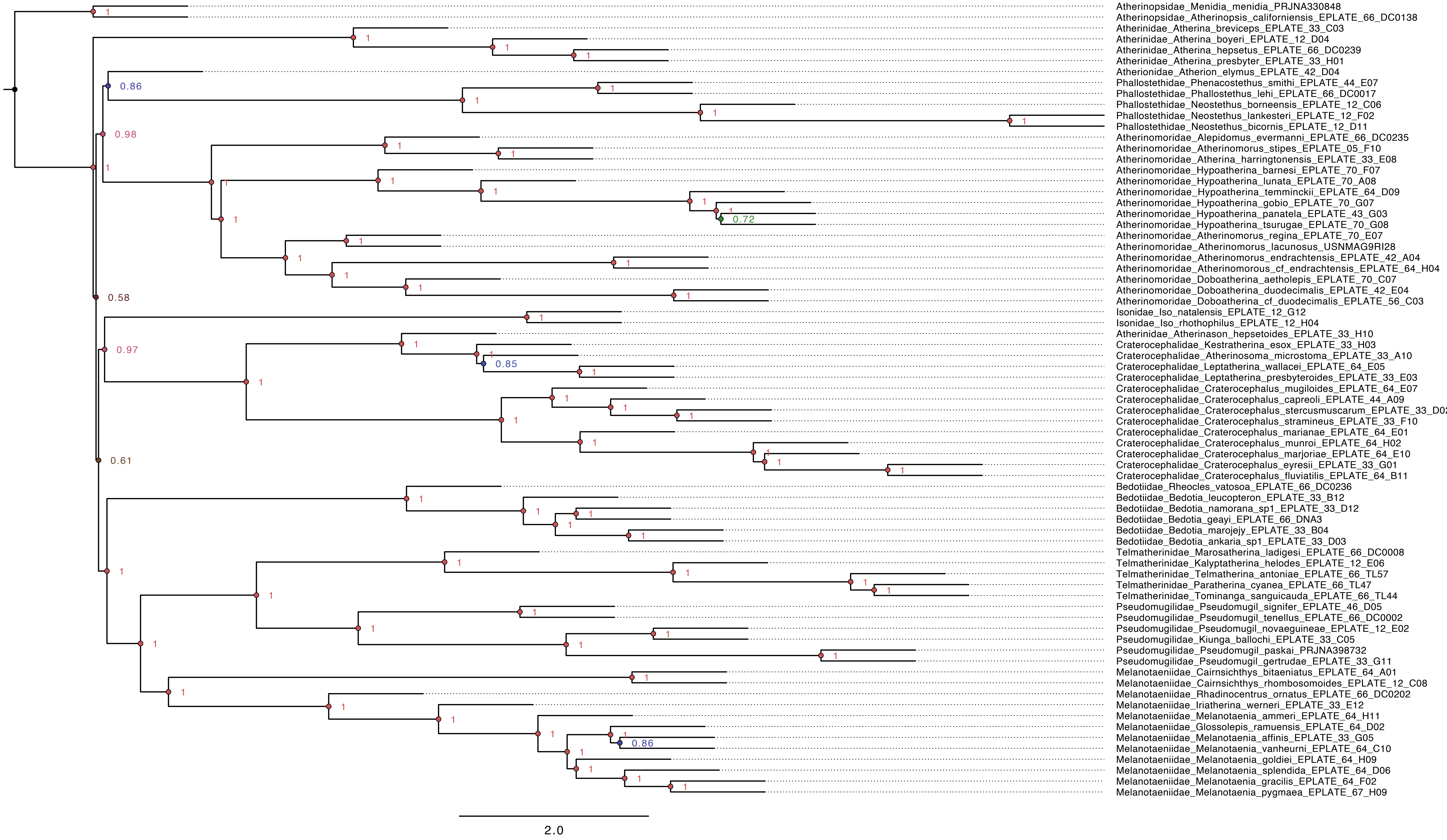

**Fig. S19: ASTRAL species tree of Atherinoidei estimated from G90 gene trees with reduced outgroups (Table 1: analysis 19). Node values indicate local posterior probability.**

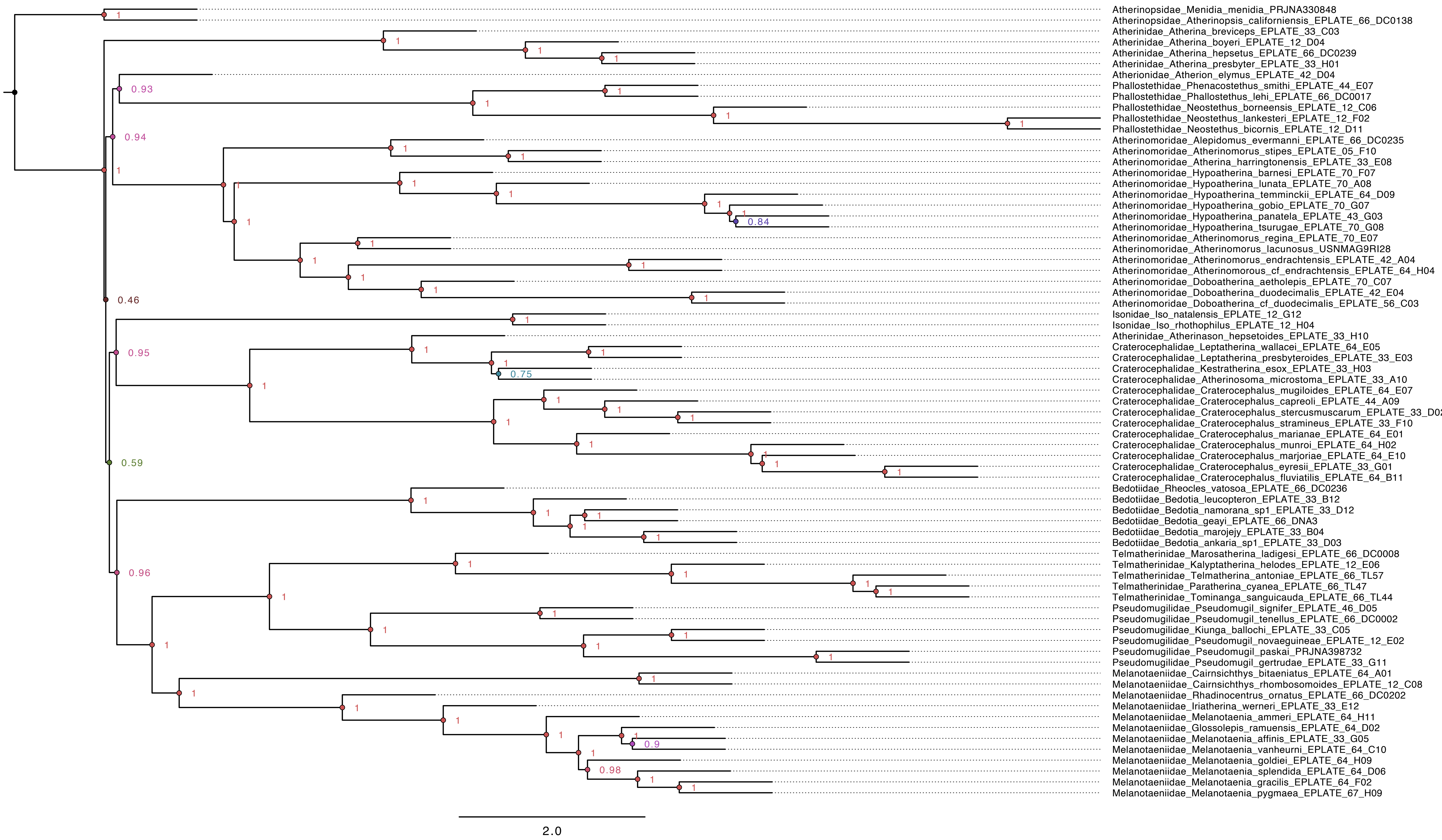

#### Fig. S20: Most likely ancestral states from DEC model on the concatenated Atherinopsidae topology.

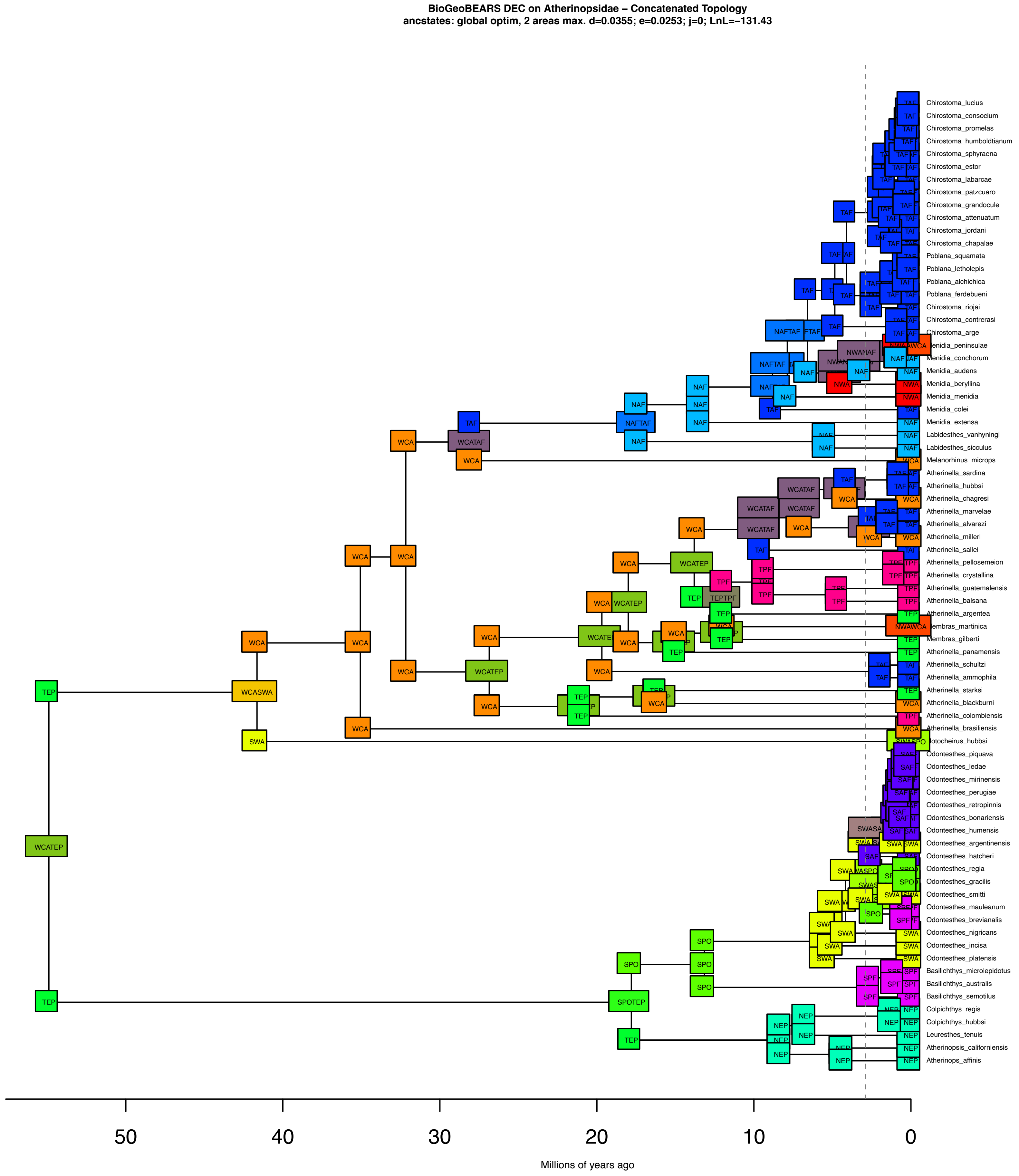

**Fig. S21: Inferred ancestral states as pie charts from the DEC model on the concatenated Atherinopsidae topology.**

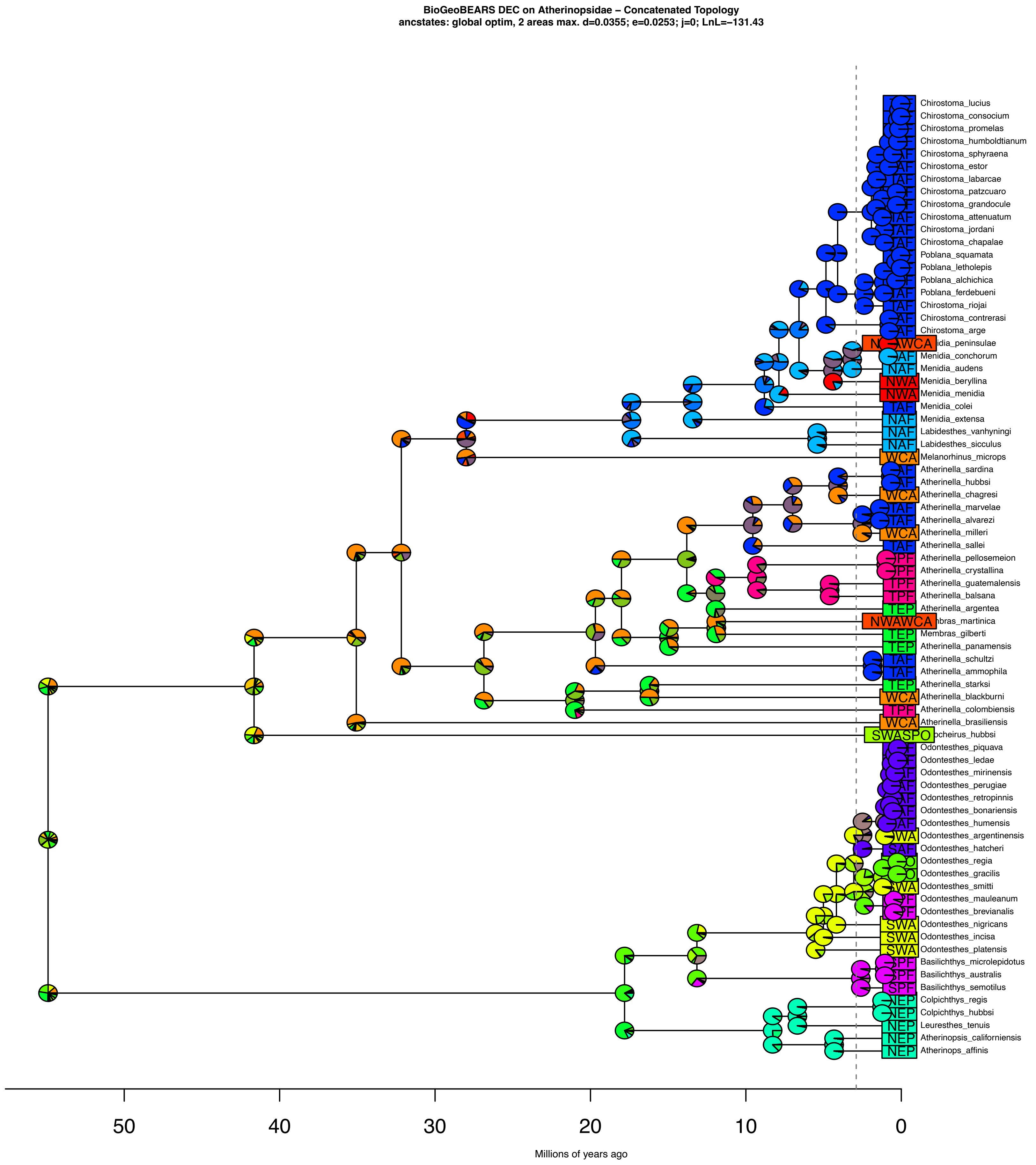

**Fig. S22: Most likely ancestral states from DEC model on the MSC Atherinopsidae topology.**

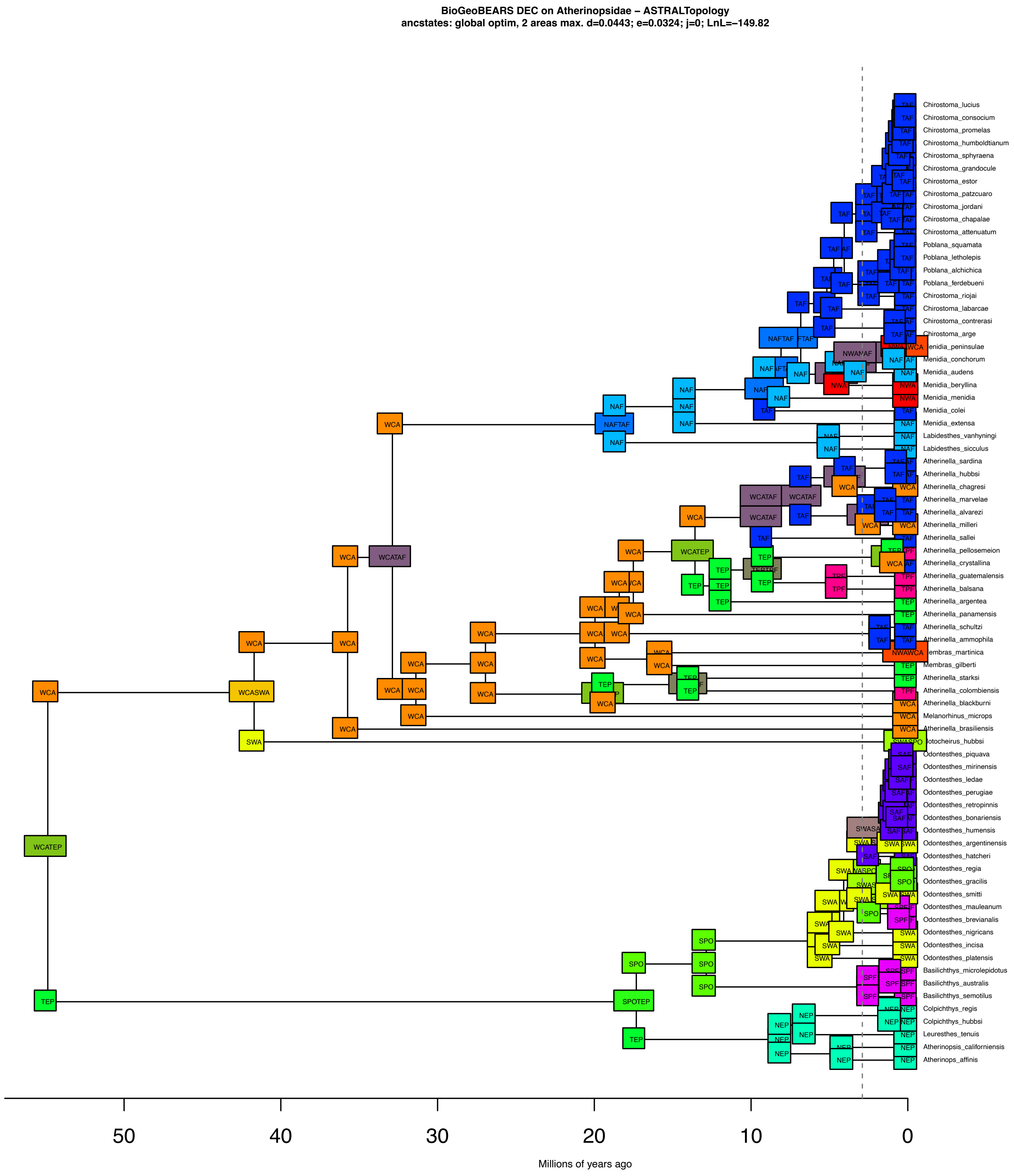

### Fig. S23: Inferred ancestral states as pie charts from DEC model on the MSC Atherinopsidae topology.

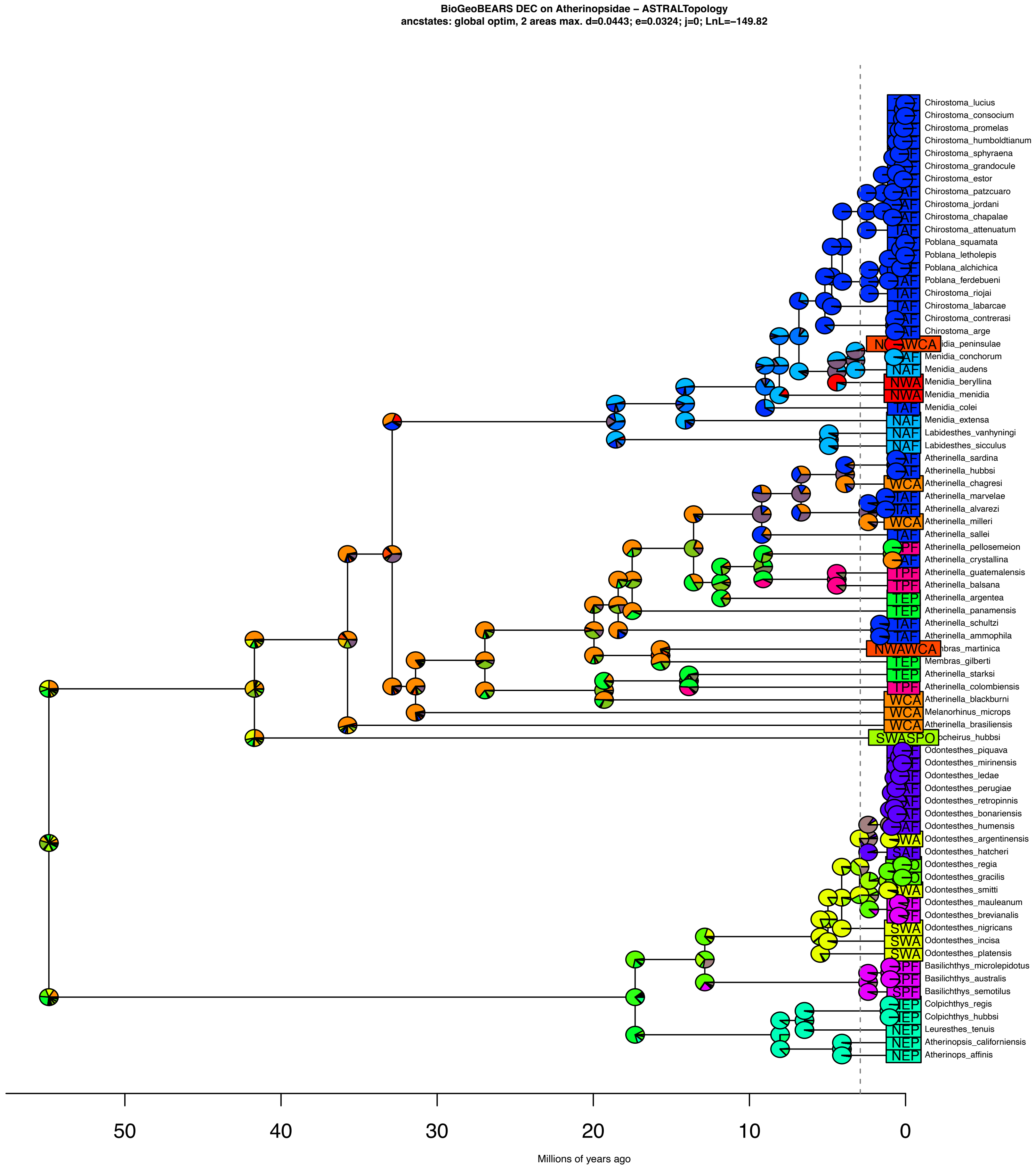

**Fig. S24: Estimated ancestral states for Atherinoidei under the DEC model using the MSC topology. Dispersal probability across the Eastern Pacific Barrier was not restricted.**

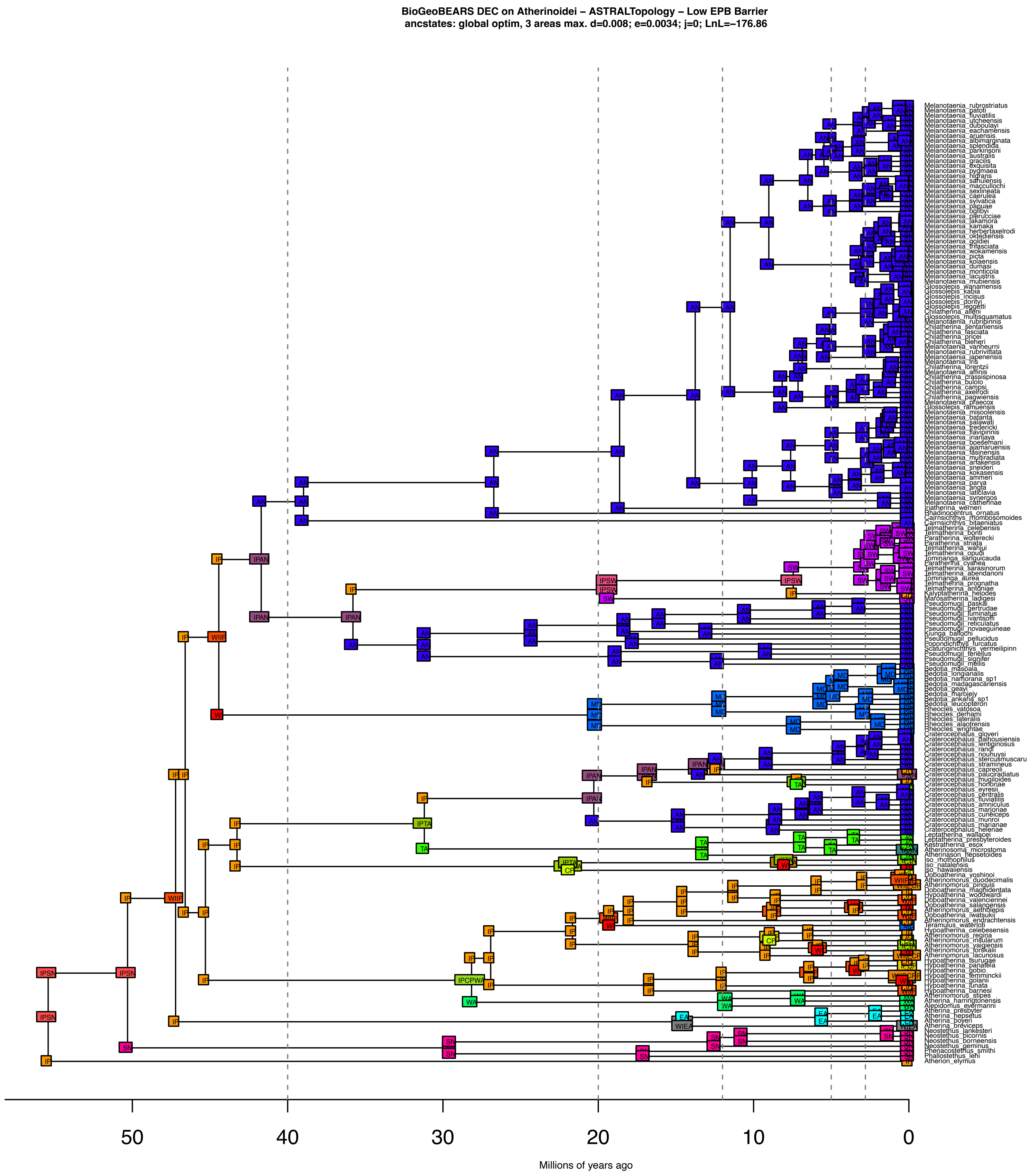

**Fig. S25: Estimated ancestral states as pies for Atherinoidei under the DEC model using the MSC topology. Dispersal probability across the Eastern Pacific Barrier was not restricted.**

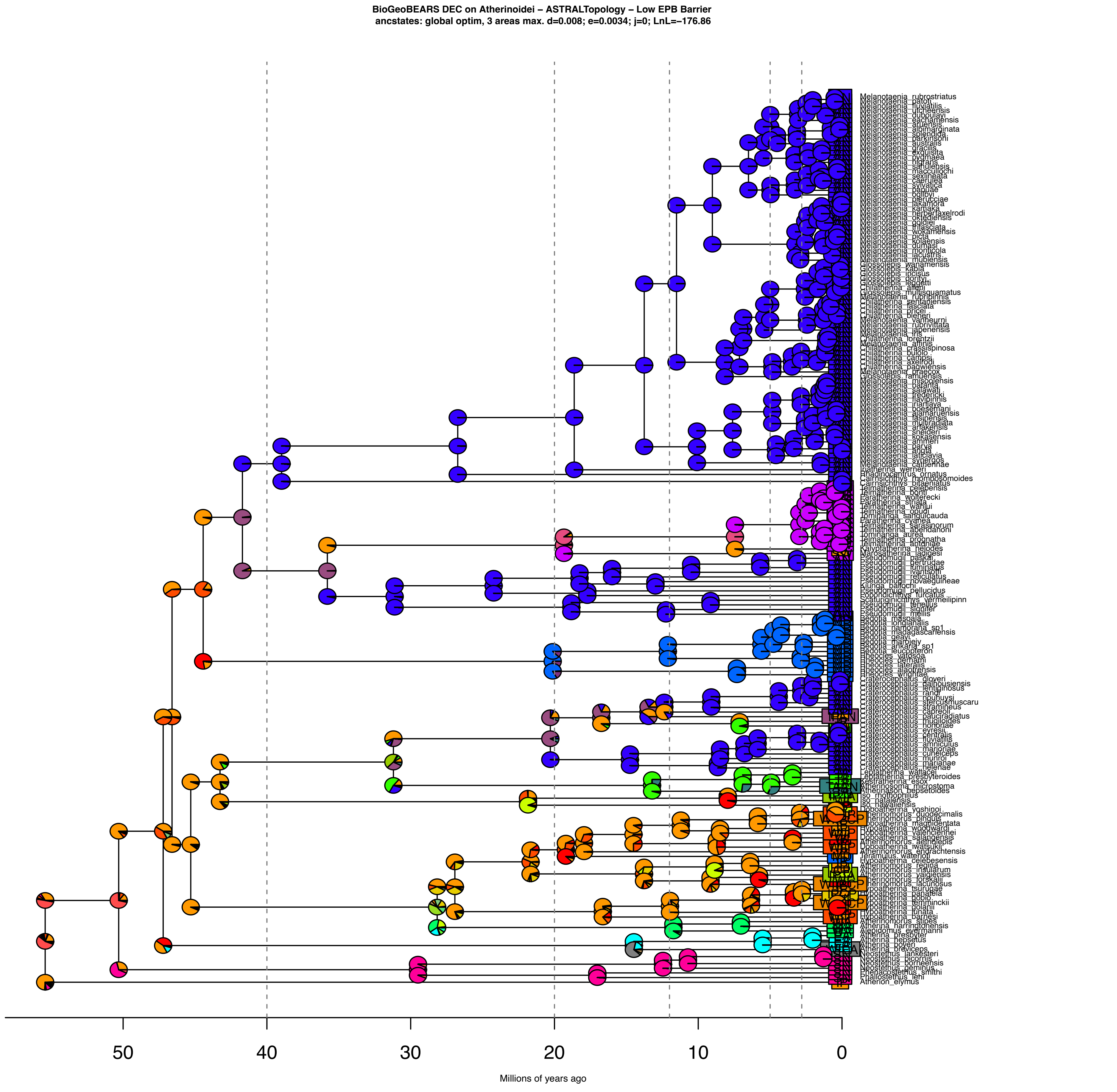

**Fig. S26: Estimated ancestral states for Atherinoidei under the DEC model using the MSC topology. Dispersal probability across the Eastern Pacific Barrier was restricted to 0.001.**

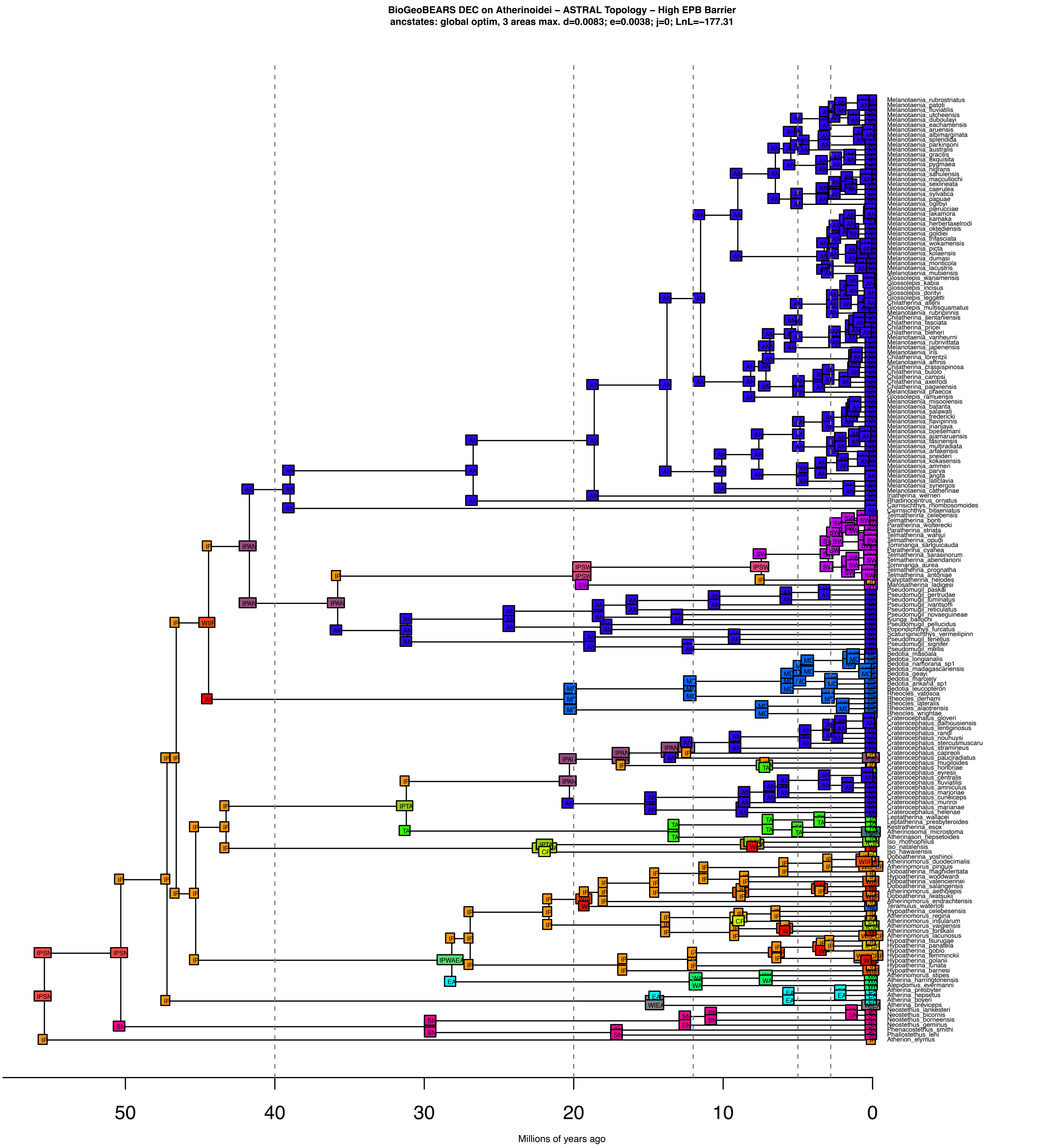

**Fig. S27: Estimated ancestral states as pies for Atherinoidei under the DEC model using the MSC topology. Dispersal probability across the Eastern Pacific Barrier was restricted to 0.001.**

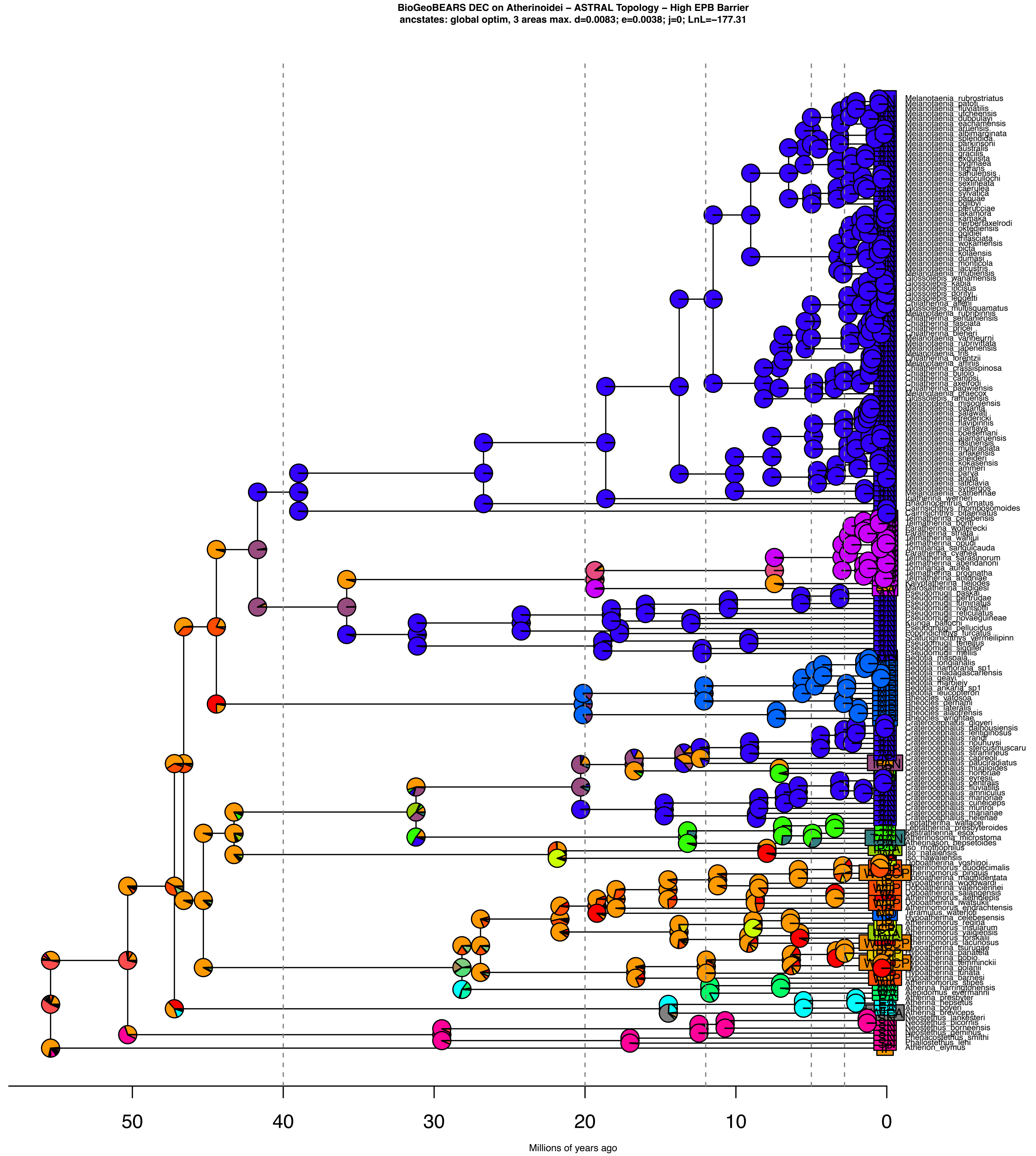

**Fig. S28: Estimated ancestral states for Atherinoidei under the DEC model using the concatenated topology. Dispersal probability across the Eastern Pacific Barrier was not restricted.**

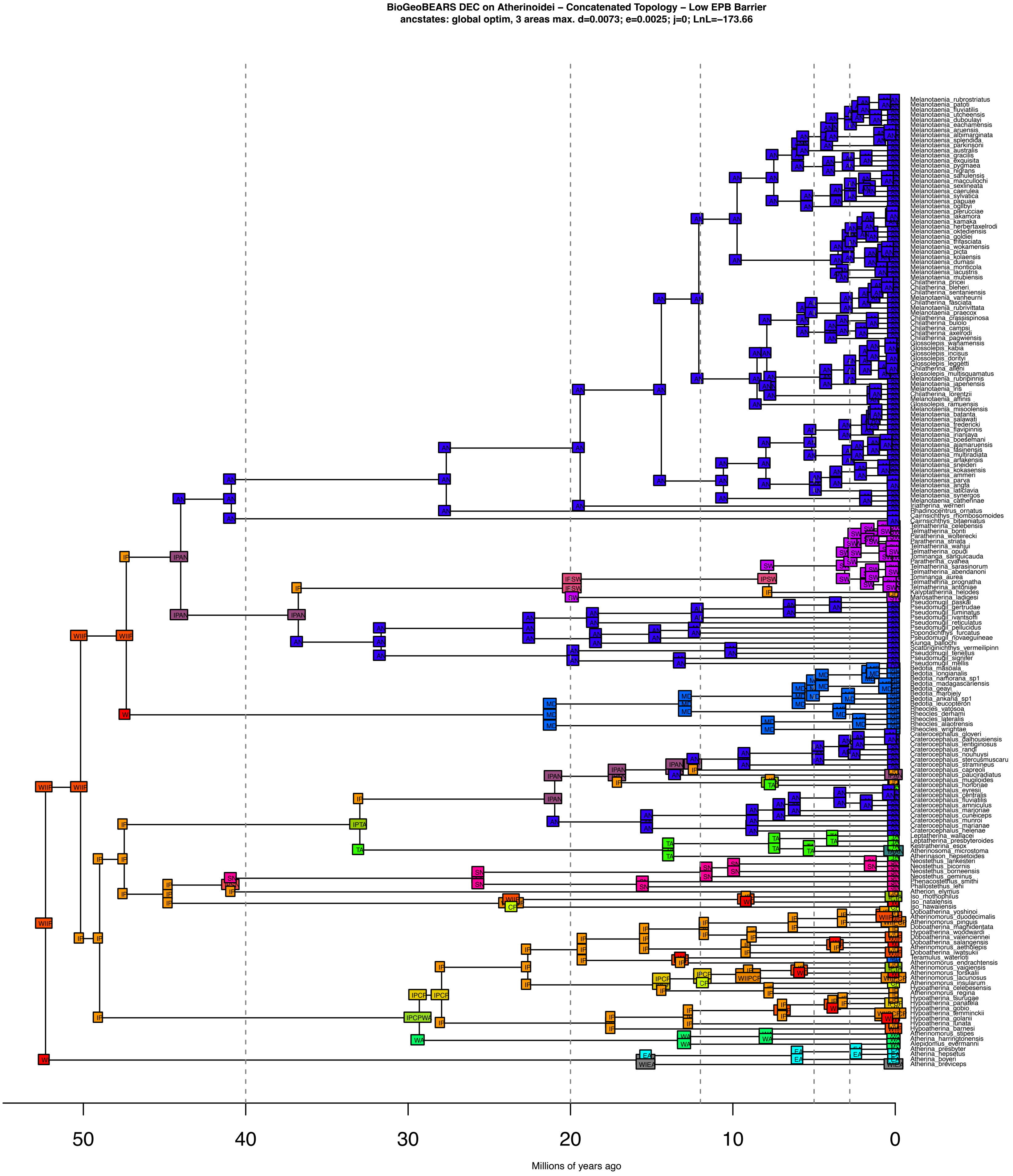

**Fig. S29: Estimated ancestral states as pies for Atherinoidei under the DEC model using the concatenated topology. Dispersal probability across the Eastern Pacific Barrier was not restricted.**

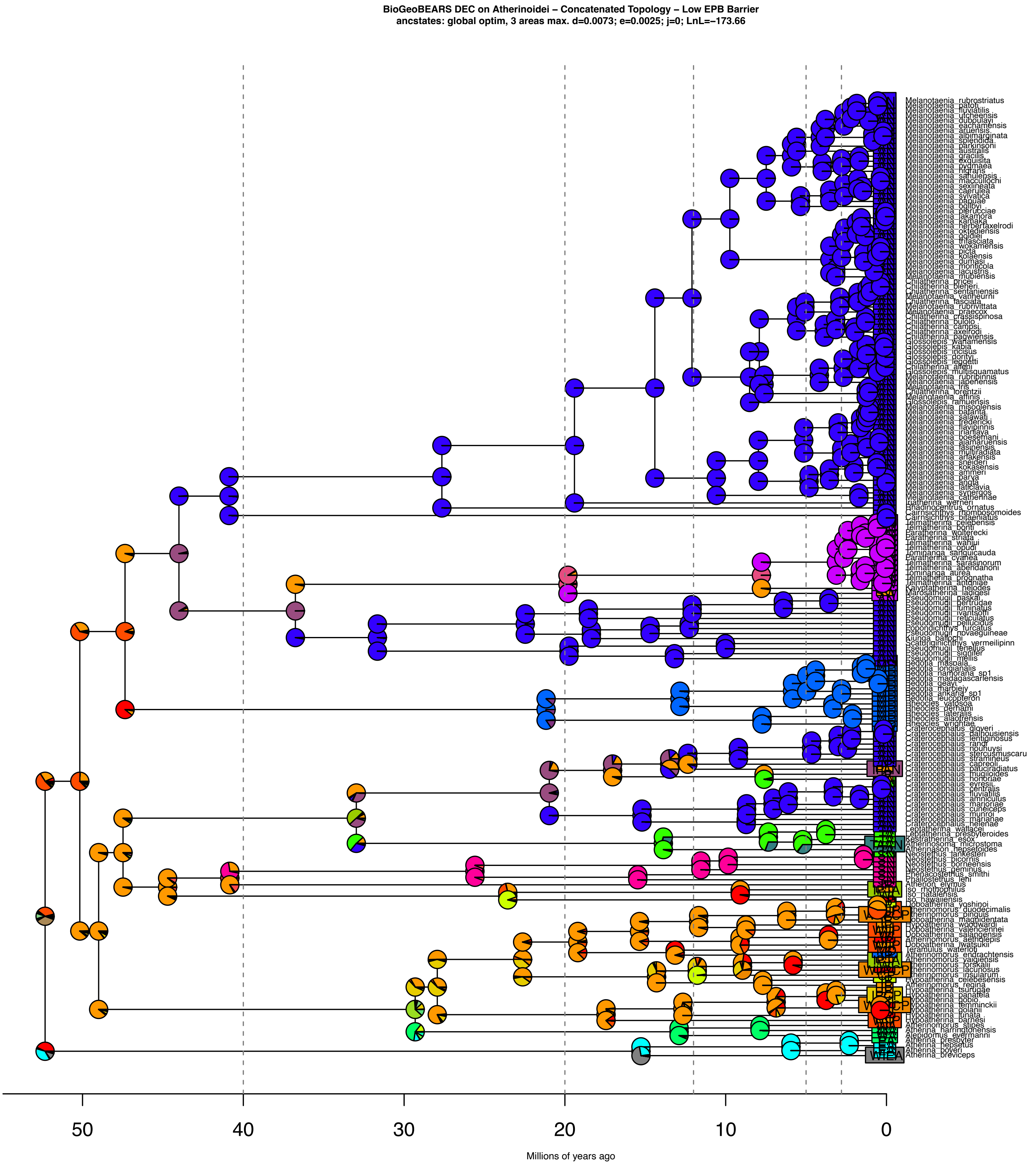

**Fig. S31: Estimated ancestral states for Atherinoidei under the DEC model using the concatenated topology. Dispersal probability across the Eastern Pacific Barrier was restricted to 0.001.**

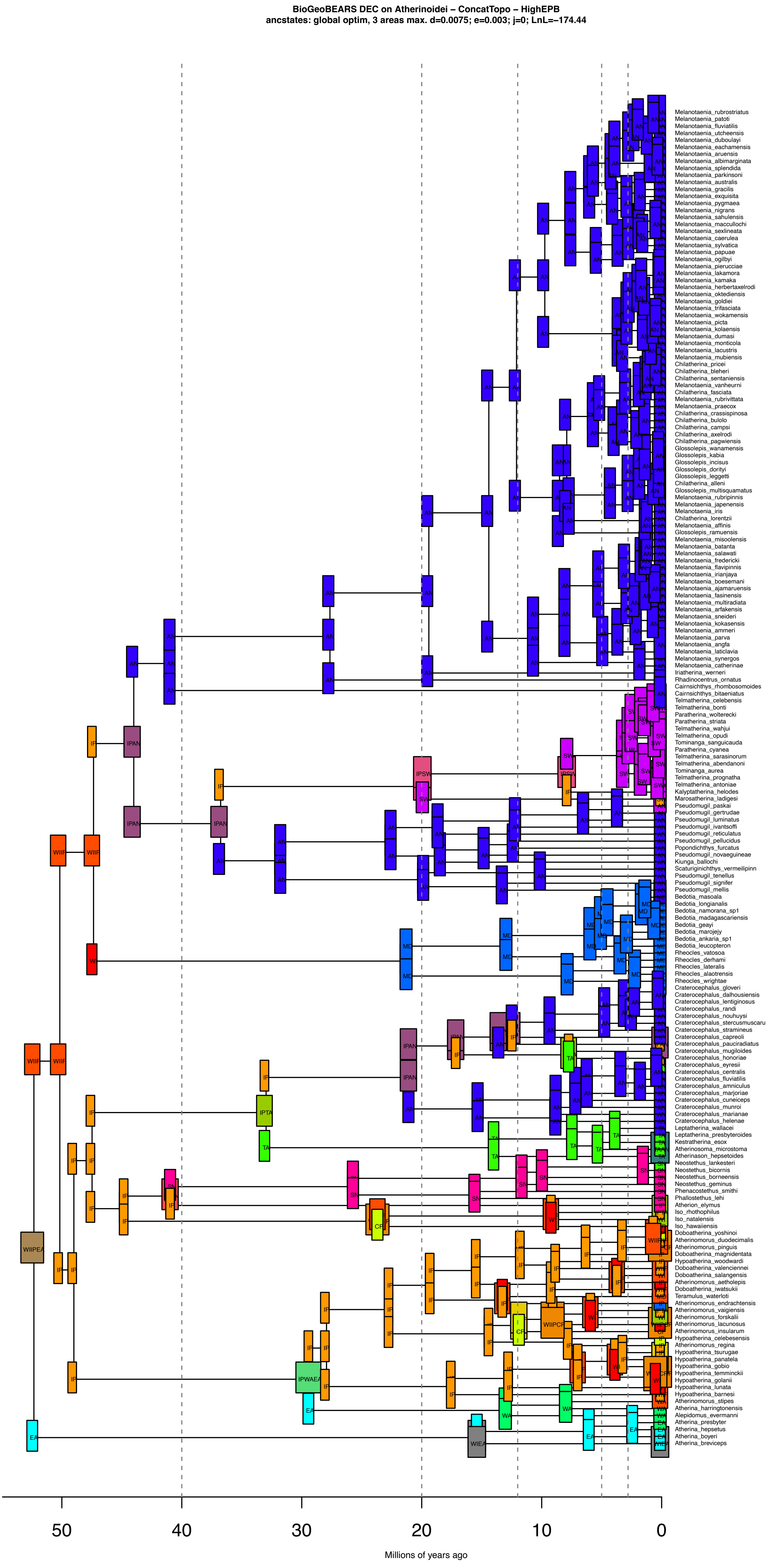

**Fig. S31: Estimated ancestral states as pies for Atherinoidei under the DEC model using the concatenated topology. Dispersal probability across the Eastern Pacific Barrier was restricted to 0.001.**

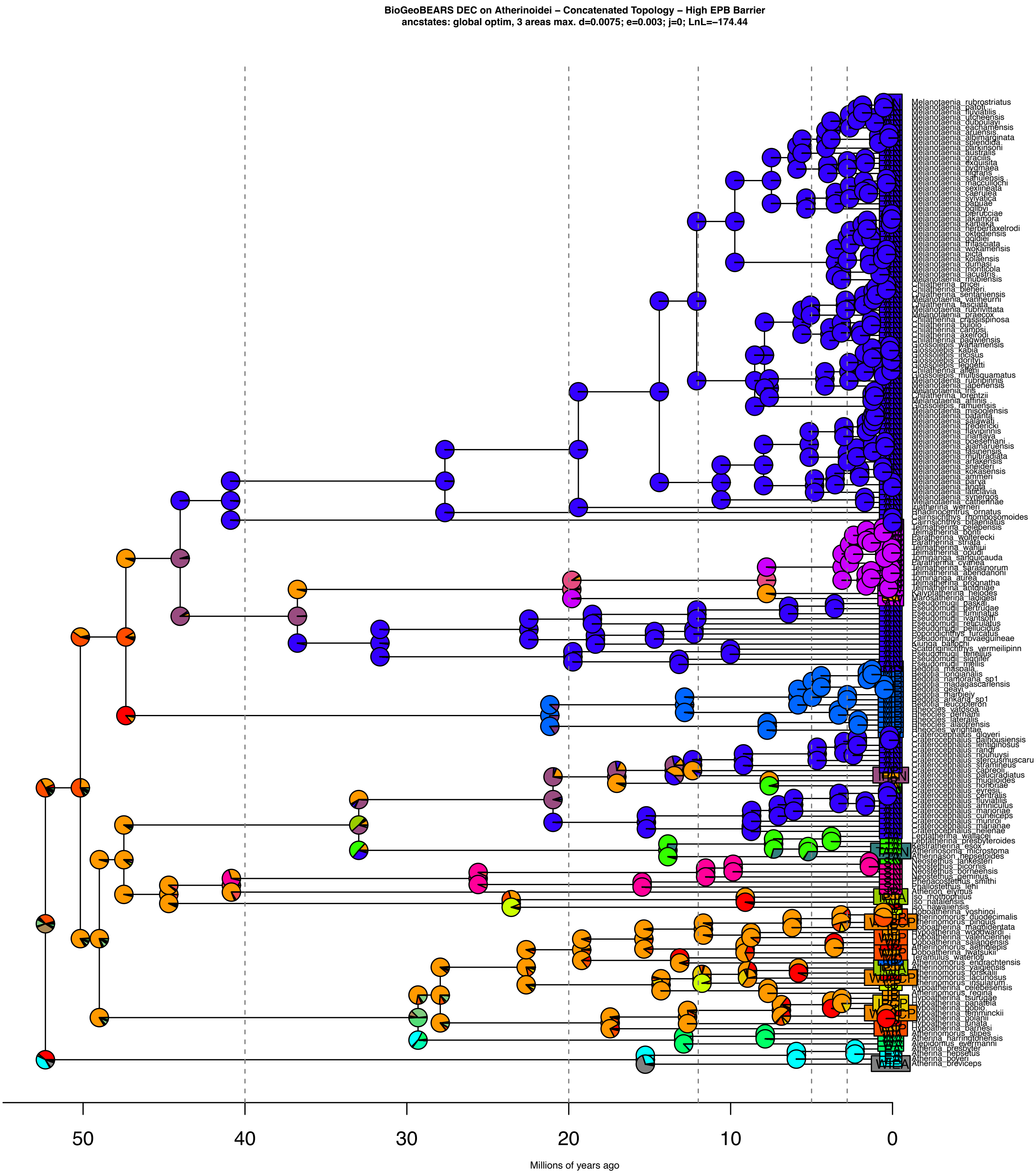
